# Sorghum *ANTHRACNOSE RESISTANCE GENE3* Is a Non-Coding RNA That Confers Fungal Resistance through Enhanced Cell Death

**DOI:** 10.64898/2026.09.18.752636

**Authors:** Demeke B. Mewa, Adedayo Adeyanju, Athanas Guzha, Pascal Okoye, Chao-Jan Liao, Gezahegn Girma, Eunyoung Seo, Michael Gribskov, Damon Lisch, Tesfaye Mengiste

**Author notes:** (Corresponding author), Mengiste, Tesfaye.

## Abstract

The fungal pathogen *Colletotrichum sublineola* is a major constraint to sorghum production. The sorghum genotype IS18760 is resistant to multiple *C. sublineola* strains, but the genetic basis of this resistance is unknown. Bulk segregant analysis using whole-genome resequencing of a biparental mapping population identified a major resistance locus, designated *ANTHRACNOSE RESISTANCE GENE LOCUS 3 (ARG3)*, associated with extensive cell death. Fine mapping delimited the *ARG3* locus to a 30 kb interval containing three predicted protein-coding genes and one unannotated non-coding RNA gene. Among these, only the non-coding RNA gene showed consistent sequence polymorphisms between resistant and susceptible lines and elevated pathogen-induced expression in the resistant parent, suggesting that it underlies *ARG3*-mediated resistance. Regions syntenic to *ARG3* across grass species contain conserved non-coding sequences enriched for H3K27me3 and H2A.Z chromatin marks, suggesting a conserved regulatory function. The resistance allele is absent from a 111-line sorghum pangenome. The resistant sorghum line harboring *ARG3* and the corresponding avirulent pathogen strain originate from the same geographic region, suggesting co-evolution between host resistance and a prevalent pathogen strain. Phylogenetic analysis suggests that loss of resistance in susceptible lines may relate to insertion of a 12.4 kb repetitive sequence. Importantly, silencing of the *ARG3* transcript in the resistant parental line abrogated resistance to *C. sublineola*, confirming that *ARG3* non-coding RNA gene underlies the fungal resistance mediated by the *ARG3* locus in IS18760. These lines of evidence show that *ARG3* is an important target for resistance breeding and provide new insight into non-coding RNA-mediated disease resistance.

**Highlight:** A rare, evolutionarily conserved allele of the non-coding RNA gene *ARG3* confers resistance to sorghum anthracnose disease by enhancing cell death at the infection site.

## Introduction

Sorghum (*Sorghum bicolor*) is a staple food in many developing nations, a major feed crop, and a potentially important source of biofuel. Anthracnose disease caused by the hemi-biotrophic fungal pathogen *Colletotricum sublineola* (Cs) reduces productivity of sorghum (Cota *et al*., 2017). The fungus exhibits significant genetic variability, making it challenging to breed for broad-spectrum and stable resistance, an issue further complicated by an incomplete understanding of the genetic mechanisms underlying anthracnose resistance (Wharton & Julian, 1996; Wharton *et al*., 2001). The genetic variation in the fungus is matched by genetic variation in resistance in the crop (Erpelding & Prom, 2004; Erpelding & Prom, 2006; Erpelding, 2008; Sharma *et al*., 2012). To explore this variation, germplasm selections for sorghum anthracnose resistance have been conducted since the 1930’s (Chen, 1934), and inheritance of the trait has been studied since 1950’s (LeBeau & Coleman, 1950). Genetic inheritance and molecular mapping of sorghum anthracnose resistance loci have been investigated for over three decades.

Genetic studies of sorghum anthracnose resistance have identified many resistance loci (Mehta *et al*., 2005; Da Costa *et al*., 2011). These studies were based on sorghum differential lines that exhibit distinct race-specific resistance to anthracnose (Cardwell *et al*., 1989; Prom *et al*., 2012) as well as the availability of a diverse germplasm collection (Casa *et al*., 2008; Cuevas *et al*., 2017; Girma *et al*., 2020). Significant progress has been made in mapping anthracnose resistance genomic regions using diverse resistance sources. Eight genomic regions were reported using association mapping in 242 diverse landrace worldwide mini-core collection, of which seven regions harbor host resistance related genes, two of which were NLR genes (Upadhyaya *et al*., 2013). Other GWAS studies also identified resistance genomic regions using diverse germplasm and multi-location trials (Birhanu *et al*., 2024). A genetic mapping study revealed two QTL that carry defense related genes (Felderhoff *et al*., 2016). A genetic mapping study of resistance to several sorghum foliar diseases identified QTL associated with plant color (Mohan *et al*., 2010). Despite this progress, identification of specific sorghum genes and proteins remain limited compared to those in other cereal crops.

Plant disease resistance is broadly categorized as either qualitative, controlled by one or a few major genes, or quantitative, involving multiple genes. Sorghum resistance to anthracnose involves both types and involves complex interactions shaped by host and the pathogen genetics. Qualitative resistance, typically mediated by simply inherited alleles, has been successfully exploited in breeding programs targeting biotrophic pathogens. A prominent example is rust resistance in wheat, where genetic resistance has been effectively used to manage a historically devastating disease (King *et al*., 2024). Resistance to certain host-specific necrotrophic pathogens is also conferred by simply inherited resistance genes or alleles that encode host-encoded proteins that detoxify specific fungal toxins, for example, the *Hm1* gene in maize, which confers resistance to *Cochliobolus carbonum* (Johal & Briggs, 1992). Resistance can also arise from recessive alleles of host susceptibility genes targeted by fungal toxins (Faris & Friesen, 2020). Such monogenic resistance genes confer highly specific resistance to certain pathogen races that produce toxins or effector proteins whether the pathogen is biotrophic or necrotrophic, although the underlying resistance mechanisms differ (Liao *et al*., 2022).

Single loci of large effect host resistance are generally mediated by effector-triggered immunity (ETI), in which intracellular nucleotide-binding leucine-rich repeat (NLR) receptors recognize pathogen effectors and activate strong defense responses, including oxidative bursts, calcium influx, immune gene activation, and hypersensitive cell death, all of which restrict pathogen growth (Jones *et al*., 2024). Structural studies further show that activated NLRs assemble into resistosomes, calcium-permeable channels that regulate ETI-associated cell death (Wang *et al*., 2019; Martin *et al*., 2020). In contrast, quantitative resistance is typically polygenic and broadly effective across pathogen races. Quantitative resistance is often associated with pattern-triggered immunity (PTI), initiated when cell-surface pattern-recognition receptors (PRRs) detect conserved pathogen molecules and activate basal immune responses (Boller & Felix, 2009; Ngou *et al*., 2022).

Plant immunity is mediated by diverse genes that encode structurally distinct proteins. PRRs and NLRs are key proteins that regulate plant immunity by mediating early steps in pathogen recognition that activate signaling cascades. Accumulating evidence implicates non-coding RNAs (ncRNAs) including microRNAs, long non-coding RNAs, and circular RNAs in plant immune functions through regulation of downstream genes (Song *et al*., 2021). In sorghum, both small RNAs and miRNAs differentially accumulate in response to anthracnose infection, in resistant and susceptible sorghum genotypes although their biological significance remains unclear (Fu *et al*., 2020). Expression levels of 50 miRNAs significantly differed between resistant and susceptible lines. Notably, the *ARG1* gene, which encodes an NLR, is located within the intron of a natural antisense RNA gene that is flanked by miniature inverted-repeat transposable elements (MITEs) that generate small RNAs (Lee *et al*., 2022). These observations suggest non-coding RNAs to be key players in plant immunity to anthracnose and other plant pathogen interactions.

Progress in identification and characterization of sorghum anthracnose resistance genes has been slow, lagging behind other pathosystems. Although many resistance QTLs have been mapped to genomic intervals containing candidate resistance genes (Cruet-Burgos *et al*., 2020), no specific resistance genes were cloned until recently. Over the past several years, we identified *ANTHRACNOSE RESISTANCE GENES 1, 2, 4,* and *5*, all of which encode NLR proteins that confer resistance to anthracnose in sorghum (Lee *et al*., 2022; Mewa *et al*., 2023; Habte *et al*., 2024). In the current report, we show that *ANTHRACNOSE RESISTANCE GENE 3* (*ARG3*), a previously unannotated non-coding RNA gene, is required for resistance to sorghum anthracnose disease caused by *C. sublineola*. *ARG3* shows consistent sequence polymorphisms between resistant and susceptible sorghum lines and is induced upon *C. sublineola* inoculation. Furthermore, suppressing expression of this non-coding RNA gene through virus-induced gene silencing rendered the resistant line susceptible to *C. sublineola*, indicating that it underlies *ARG3*-mediated resistance in sorghum line IS18760. Together, these results demonstrate that *ARG3* is a rare allele of a non-coding RNA gene that mediates anthracnose resistance through a mechanism involving extensive cell death that restricts fungal growth.

## Materials and Methods

### Experimental materials and disease evaluation

This study started with evaluation of sorghum lines that showed differential responses to strains of *C. sublineolum* (Prom *et al*., 2012; Mewa *et al*., 2023). The fungal strains, inoculum preparation, disease assay, and evaluation of resistance responses were as previously described (Mewa *et al*., 2023). Briefly, five strains, Csgl1, Csgl2, Csgrg, Cs27, and Cs29 were grown in petri-dishes on a lab bench under continuous light. Conidia were harvested from actively growing cultures, resuspended in water (1×10^6^ conidia/ml), and sprayed on month-old seedlings until the entire plant surface became uniformly wet. Inoculated plants were kept in a humidity chamber (>80 relative humidity) for 48 h and then transferred to greenhouse with a temperature setting of 28-32°C. An overhead mist was set to run every 30 min during the daytime until the disease fully developed in susceptible plants, a week to 10-days after inoculation. Resistant and susceptible plant genotypes were primarily identified by visual observation. Evaluation of host responses was further examined using a detached-leaf disease assay and trypan blue staining and PCR quantification of the fungal growth on the host tissue.

### Mapping population and molecular data

We identified a line, IS18760, that is resistant to two of the five known strains of *C. sublineola* (Cs27 and Cs29). This line was crossed to the highly susceptible line TAM428, and a biparental mapping population was developed. F2 plants that showed the parental-type host responses to Cs29 were advanced to the F5 generation, with multiple plants from each lineage being tested for consistent responses prior to advancement. In every disease assay, the parental lines were included as controls. At F5, 50 resistant plants and 50 susceptible plants, each from a unique F2 plant, were used for bulk-segregant analysis. High-quality genomic DNA was extracted from each F5 plant. Equal amounts of DNA from each sample were pooled to make a resistant bulk sample and a susceptible bulk sample. The two bulk samples and the genomic DNA of IS18760 were resequenced separately at 25X coverage using the Illumina NovaSeq 6000 platform. Genomic DNA sequence data from TAM428 was available from a previous study (Lee *et al*., 2022).

### *ARG3* mapping and characterization

Bulk-segregant analysis of genomic DNA sequence data (BSA-seq) was used to map the genomic regions that harbor the resistance loci based on BTx623 V3.1.1 and Rio V2 sorghum reference genomes using the QTL-seq analytical pipeline (Takagi et al. 2013). Briefly, the method utilizes single nucleotide polymorphisms (SNPs) generated from genomic raw-reads that are aligned to the reference genome. Mapping was carried out using each parental genome raw-reads as the internal reference to estimate the SNP frequencies in the bulk samples. When genomic regions are unlinked to a given phenotype, recombination is expected to occur randomly, and thus SNP-index approximates 0.5 and the difference, delta SNP-index [Δ(SNP-index)], between the bulk samples approximates zero. In contrast, in the genomic regions that are tightly linked to the phenotype, the SNP-index will approximate zero in one bulk sample and one in the other bulk sample, and Δ(SNP-index) results in a QTL peak. BSA-seq uses the average Δ(SNP-index) over 2 Mb (and again over 4 Mb) genomic windows that slide 50 kb at a time.

Fine mapping was carried out primarily using recombination analysis-based on indel size-markers that differentiate the parental lines. To identify potential indel sites in the QTL interval, the binary-alignment map (BAM) files of the parental lines and bulk-progeny sample genomes were examined on Integrated Genome Viewer (IGV). Primers flanking indel sites were designed to amplify polymorphic size markers and tested for size differences using PCR and gel electrophoresis in the parental lines before applying to plants in the mapping population. Plants with known responses were examined for phenotype to marker association to exclude genomic regions that are unlikely to harbor the causal resistance locus. A series of four marker development and recombination analyses were carried out on 109 F5 plants using a total of 15 markers. The markers in the first round of recombination analysis were sparsely distributed along the 3.8 Mb region of *ARG3* QTL with adjacent markers several hundred kilo-bases (kb) apart. Markers in subsequent recombination analyses were fewer in number, and progressively, the *ARG3* genomic interval was narrowed.

A single F5 plant that was heterozygote at the *ARG3* genomic region was identified using indel marker 55.11 (Fig. 4) and advanced to F6. Similarly, homozygote F6 plants were advanced to F7 plant families. In eight F7 resistant plant families, and in eight susceptible plant families, the true-to-type genotypes were confirmed in five or more plants per family, and all progenies were evaluated for host responses, alongside the parental lines.

The final genomic interval carrying *ARG3* locus was evaluated for presence of candidate genes based on functional annotation, DNA sequence polymorphism, and gene expression. We examined the genomic sequence of the parental lines, the bulk samples, and diverse sorghum lines using publicly available data. A transcript in the *ARG3* genomic interval that lacks an annotated gene was amplified and then sequenced using the WideSeq DNA sequencing platform by the Purdue University Core Genomics facility. qRT-PCR was used to generate relative transcript level data. Coding Potential Calculator (CPC2; https://cpc2.gao-lab.org/) was used to evaluate the coding potential of *ARG3*. Genome assembly of the parental lines, the bulk samples, and many other re-sequenced sorghum variants were carried out to examine insertion/deletion (indel) sites using the genome assembler SPAdes 3.14.1 (Prjibelski *et al*., 2020). The assembled contigs were examined using Bandage software (Wick *et al*., 2015).

For pangenome analyses, genomic sequences of *ARG3* including three major indel regions were manually extracted from Sorghumbase.gov and Phytozome plant genome database. DNA sequence alignment was carried out using Job dispatcher software; in Multiple Sequence Alignment main menu, *Kalign* with Pearson/FASTA format and *Mview* functions were used for sequence alignment. Phylogenetic analysis was carried out based on the 1,069 bp *ARG3* transcript sequence from the resistant parental line IS18760, which is identical to its genomic sequence. Tassel software was used for cluster analysis based on UPGMA function, and the pairwise distance matrix was exported as *Newick* file format. Phylogenetic tree was created using FigTree software. Synteny of the *ARG3* QTL region and comparative genomics and histone regulation of *ARG3* locus were examined using the Comparative Genomics (CoGe) (https://genomevolution.org/coge/), the plant epigenome browser (https://epigenome.genetics.uga.edu/PlantEpigenome/index.html) (Lu *et al*., 2019) and the National Center for Biotechnology Information (NCBI) platforms.

### Virus induced gene suppression assay for ARG3 suppression

Modified pCAMBIA1380-FoMV vectors (Mei *et al*., 2019) generously provided by Dr. Saet-Byul Kim (University of Nebraska) were used to clone a 253bp segment of the non-coding RNA gene (*ARG3*) from the resistant sorghum line IS18760. The constructs were generated by amplifying a 253 bp sequence from cDNA derived from the resistant sorghum line IS18760 using primers listed in Table S3 and ligated into the modified FoMV plasmid. The constructs were confirmed by sequencing before transformation into Agrobacterium. A construct containing a 300 bp fragment of sorghum ubiquitin (Sobic.010G239500) was transformed into agrobacteria for use as a positive control for suppression of gene expression because it causes small necrotic spots which is now used as a visible marker for onset of gene suppression (Singh *et al*., 2018). Four weeks old *N. benthamiana* plants were infiltrated with Agrobacterium containing the different constructs (Empty vector, Ubiquitin-VIGS and ARG3-VIGS) using three plants per construct. The infiltrated region in the leaf was marked with a permanent marker. At five days post infiltration the infiltrated regions for each construct were collected by cutting out with scissors while avoiding the veins. The leaf material was ground using a mortar and pestle in 10 mL of ice-cold phosphate-buffered saline (PBS), pH 7.2. The resulting slurry was filtered through cheese cloth into a 50 mL falcon tube. The filtrate was then poured into a petri dish before approximately 500 mg of silicon carbide (Sigma Aldrich Ref# 378097) was added and mixed thoroughly. Sorghum seedlings at the two-leaf stage were then rub inoculated by gently pinching the leaves between the thumb and index figure and gently rubbing with the mixture of the filtrate and silicon carbide. A minimum of three plants per construct were treated for the silencing experiment. This process was repeated seven days later to enhance suppression of target gene expression. The inoculated plants were grown at 25 °C under 12-hour light and dark cycles before inoculation with *C. sublineola*.

The *C. sublineola* inoculation for the disease assay was conducted at the onset of the first signs of leaf necrosis associated with the control ubiquitin silenced lines at approximately 10 days after the initial VIGS treatment. The *C. sublineola* spray inoculated plants were kept under high humidity at 25 °C, 12-hour light and dark cycles. At 5-7 days after disease inoculation the plants were sampled for quantification of fungal DNA and gene expression analysis using qRT-PCR as described earlier and diseases symptoms were visually inspected.

## Results

### Characterization of IS18760 for responses to strains of *Colletotrichum sublineola*

In a screen of sorghum germplasm for resistance to *Colletotrichum sublineola* strains, we previously found that the sorghum line IS18760 is resistant to the *C. sublineola* strains Cs27 and Cs29 but highly susceptible to strains Csgl1, Csgl2, and Csgrg (Mewa, 2020; Mewa *et al*., 2023). IS18760 exhibited resistance to high-density spore spray and drop inoculation by Cs27 and Cs29 (Fig. 1a, b). Cs27 and Cs29 were collected from Ethiopia while the other three strains are all from the U.S. Cs27 exhibited uneven growth on artificial growth media and its conidiation was poor, whereas Cs29 exhibited a steady growth and sporulated profusely, producing abundant conidia for inoculum. For this reason, Cs29 was used in all subsequent evaluations of host responses. TAM428 showed characteristic anthracnose disease susceptibility symptoms with visible signs of the pathogen deposition, whereas IS18760 showed no disease symptoms and highly restricted disease lesion that failed to expand after drop inoculation (Fig. 1a). This study was initiated to identify and characterize the loci underlying the resistance response in IS18760 to strain Cs29.

**Fig. 1.**
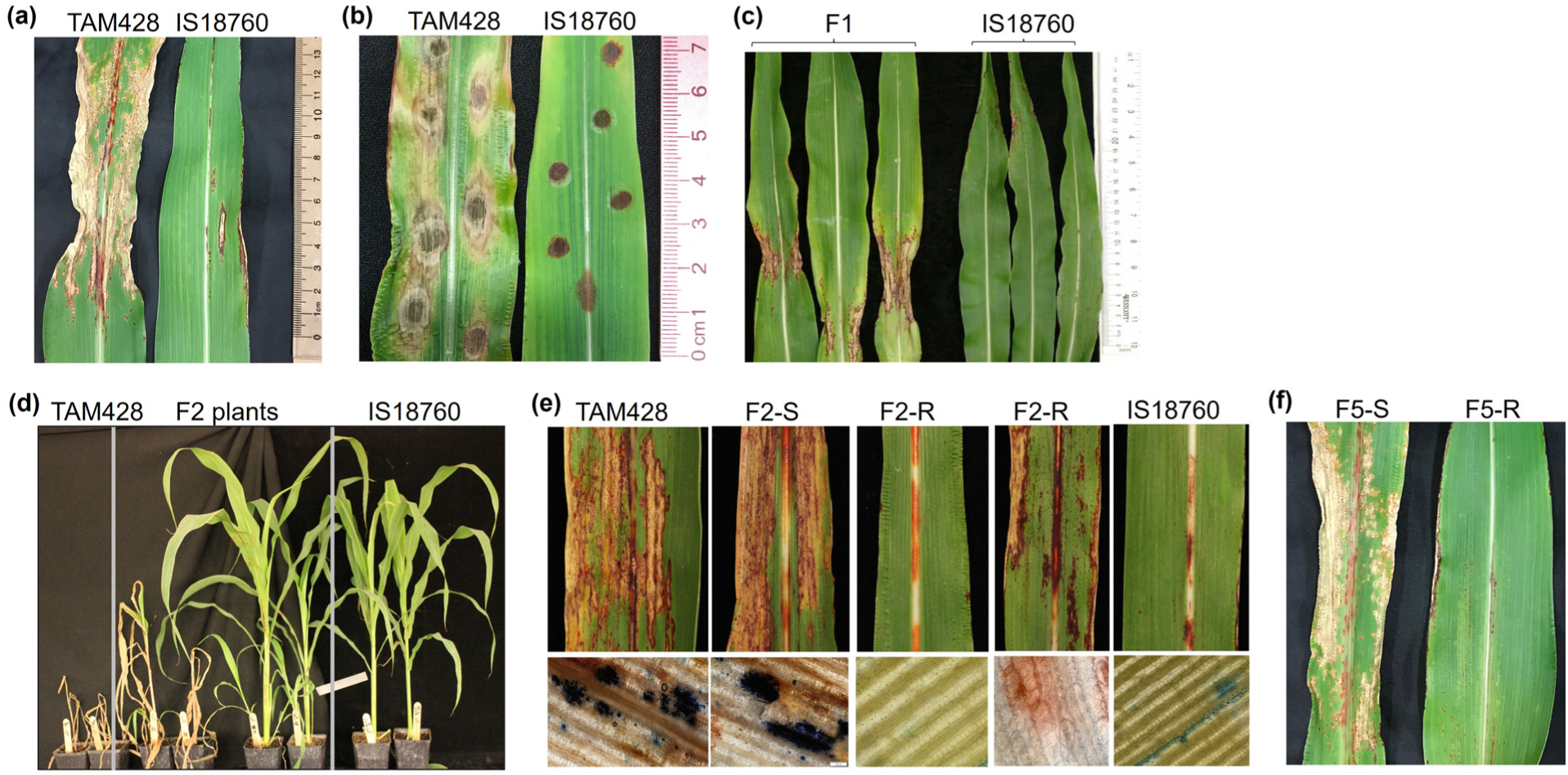
Resistance response to *Colletotrichum sublineola* in parental lines and progenies of TAM428 x IS18760 mapping population. (a) Parental lines after spray inoculation. (b) Parental lines after drop-inoculation. (c) F1 plants and the resistant parent line. (d) Whole plant view of parental lines and F2 plants. (e) F₂ plants and parental lines. F2-S = susceptible F2 plant. F2-R = resistant F2 plant. Lower panel is trypan blue staining showing fungal growth. (f) F5 sibling plants. All images were taken at 10 days post-inoculation with *C. sublineola* strain Cs29.

### Genetic mapping of the anthracnose resistance loci in IS18760

To map the genomic region underlying resistance to Cs29 in IS18760, a biparental mapping population was developed by crossing IS18760 to the highly susceptible line TAM428. F1 plants showed resistance response with extensive cell death phenotype that lacks pathogen growth (Fig. 1c). At F2, plants segregated for resistance responses observed in IS18760 and plants with susceptible response observed in TAM428 (Fig. 1d,e). Each F2 plant with clear resistance or susceptibility was advanced to the F5 generation, after disease rating at each generation. All the susceptible plants showed consistent responses in the subsequent generations, whereas the resistant plants segregated for the disease phenotype. Sample host responses of F5 plants are shown in Fig. 1f, which are typical of the disease phenotypes for advancing plants at each generation.

Bulk-segregant analysis of whole genome sequence data (BSA-seq) was carried out to identify the genomic regions responsible for the resistance in IS18760. At the F5 generation, a resistant bulk DNA sample from 50 resistant plants and a separate susceptible bulk DNA sample from 50 susceptible (Fig. 1f) plants were used for bulk-segregant analysis (BSA). BSA-Seq based on BTx623 and the Rio reference genomes revealed consistent QTL results. A single highly significant (p-value <0.01) QTL was identified towards the distal end of chromosome 04 in which the causal locus was designated as *ANTHRACNOSE RESISTANCE GENE3* [*ARG3*)] (Fig. 2). The QTL was identified based on polymorphisms in 2 Mb sliding windows using the well refined BTx623 reference genome (Table S1). Like TAM428, BTx623 is susceptible to Cs29, and they exhibited high sequence similarity in this QTL region. The ΔSNP-index estimate, which is the basis for QTL test of significance, ranged from 0.31-0.64 with a 0.30 threshold at 95% confidence interval (CI) and a 0.39 threshold at 99% CI. The genomic interval for the 95% CI was 53.50-60.20 Mb and the interval for the 99% CI was 52.50-61.85 Mb. In the wider 9.4 Mb interval at 99% CI, the read-depths were 32-40 (average=37) and the SNP counts were 1,171-2,904 (average=1,922).

**Fig. 2.**
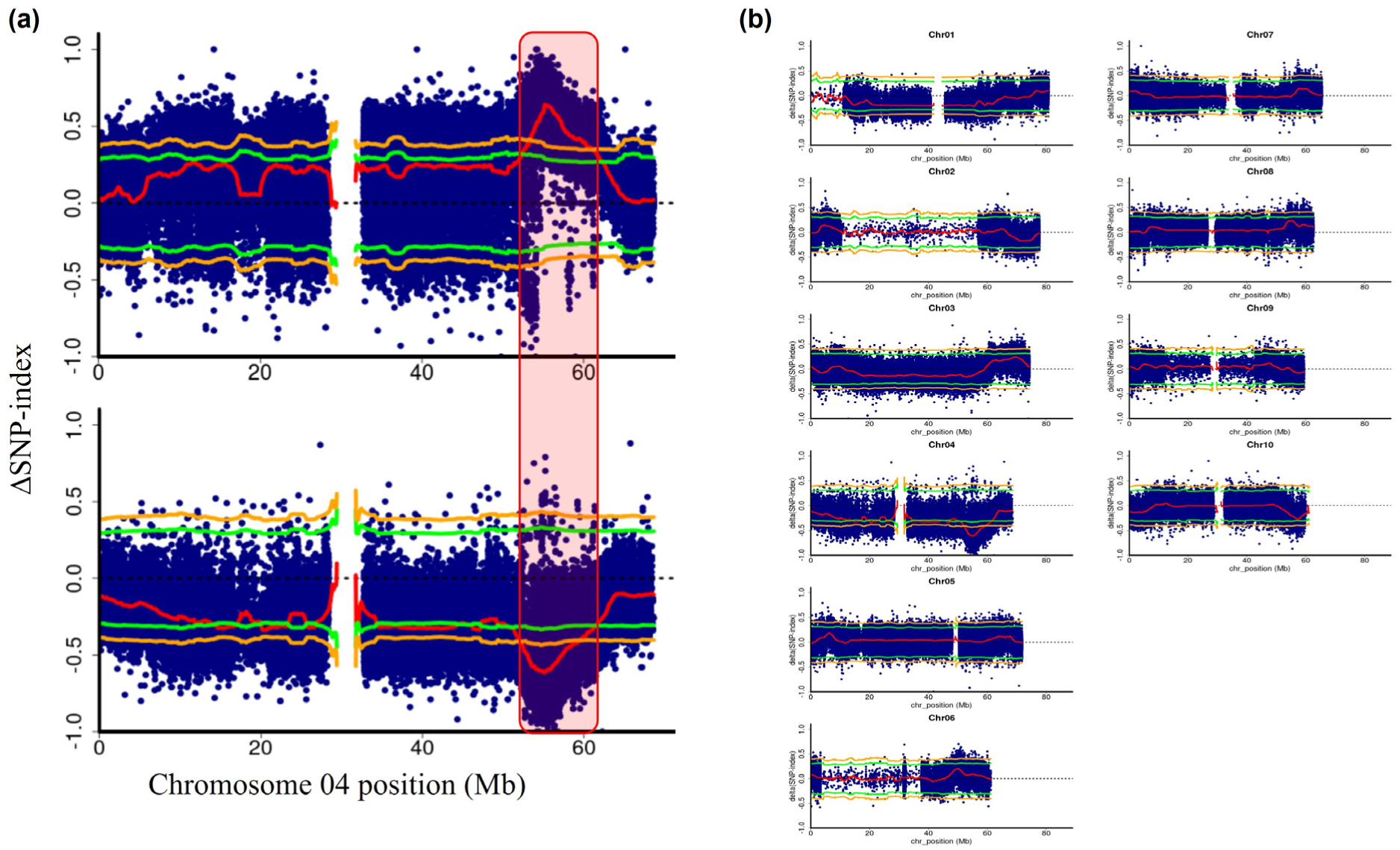
Sorghum anthracnose resistance locus identified through BSA-Seq. (a) The single highly significant QTL map on chromosome 4. The red line shows the average ΔSNP-index across the sliding window, and the green and orange lines show statistical thresholds (*p-value = 0.05 and 0.01*). The upper panel was generated with respect to the genome sequence of the resistant parental line and the lower map with respect to the susceptible parental line. (b) QTL map across the ten chromosomes of the sorghum reference genome BTx623 v3.1.1.

### ARG3-mediated resistance in NIL lines exhibited extensive cell death

The *ARG3* phenotype was examined using F7 families that shared a large proportion of the genome but varied at the *ARG3* locus region. These F7 families were developed from a single resistant F5 that was a heterozygote at the *ARG3* locus (*ARG3/arg3*). F7 nearly isogenic plant families carrying the true-to-type parental genotypes (*ARG3/ARG3* and *arg3/arg3*) were evaluated for disease phenotype and fungal growth (Fig. 3). We reasoned that these plant families had similar genomic backgrounds, and that the small proportion of variable genomic regions were randomly distributed in the resistant and the susceptible plant families. Thus, the overall difference in the resistance response can be attributed to *ARG3* locus. Plants that carried the susceptible genotype showed chlorosis and extensive necrotic lesions, with pathogen structures clearly visible, whereas plants that carried the resistant genotype showed no chlorosis and inhibited pathogen growth (Fig. 3a,b,c). In contrast to the resistant parent (Fig. 1a,b), F1 plants and resistant plants in the F7 population showed circular and elliptical spots of dead tissue and extensive cell death suggesting that the extensive cell death is attributable to ARG3-mediated resistance response (Fig. 1c, Fig. 3). The presence of both restricted and extensive cell death phenotypes in the resistant F7 category suggests that the F5 plant used to develop these nearly isogenic lines segregated for other QTL that modify the quality of resistance in these lines. Trypan blue staining and qPCR quantification of the fungal DNA showed a clear-cut difference in support of fungal growth in the susceptible parental lines and these nearly isogenic lines (Fig. 3b,c). Collectively, our analyses suggests that the wide-spread cell death that prohibits pathogen proliferation is mediated by the *ARG3* locus. This differs from the canonical effector triggered immunity (ETI) hypersensitive response and that similar resistant phenotype of the resistant parent.

**Fig. 3.**
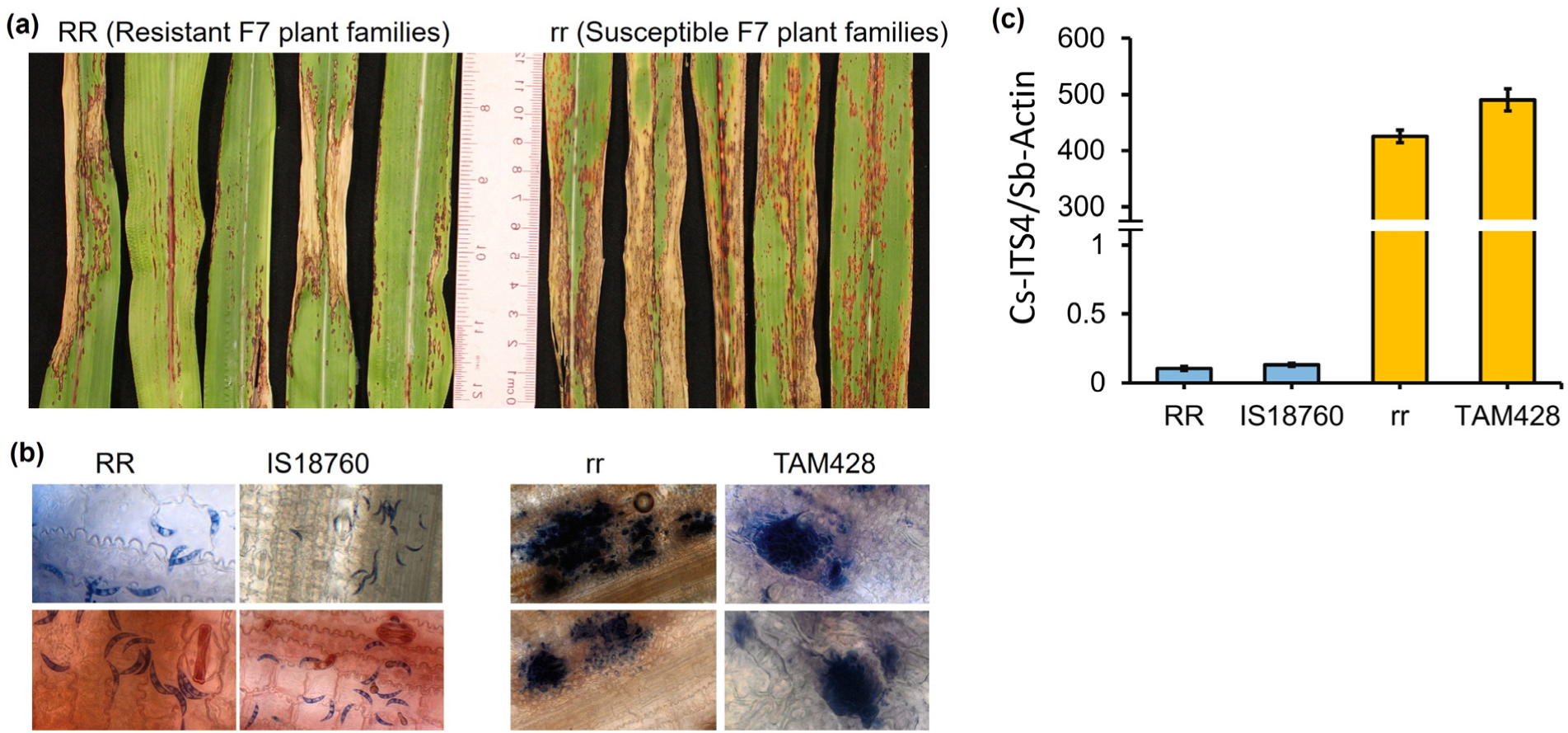
The *ARG3* disease response phenotype and restricted fungal growth in resistant genotypes. RR (*ARG3/ARG3*, resistant) and rr (*arg3/arg3*, susceptible) genotypes are F7 nearly isogenic lines. (a) Representative resistance phenotypes in eight resistant plant families and eight susceptible plant families. (b) Trypan blue staining to reveal fungal growth on host tissue of nearly isogenic and parental lines. (c) Relative quantification (qPCR) of the fungal growth using *C. sublineola ITS4* gene (Cs-ITS4) relative to sorghum Actin gene (Sb-Actin) seven days after inoculation.

### Fine mapping of the *ARG3* locus

Fine mapping of the *ARG3* locus was carried out based on analysis of recombination frequencies between markers in the F2 population. Whereas the susceptible F2 plants showed true-to-type susceptible response in subsequent generations, some of the resistant progenies segregated suggesting that resistance was dominant over susceptibility. Phenotype to marker-genotype association in the recombination analyses also supported the hypothesis that the allele in the resistant parental line was dominant over the allele in the susceptible parental line, and thus the heterozygote genotype was resistant (Table S2), which in turn facilitated the fine mapping.

Recombination analysis was carried out of a 3.8 Mb *ARG3* genomic region carrying the highest ΔSNP-index estimates across the 50 kb sliding window using 109 F5 plants (Fig. 4a, Table S2). PCR-based genotyping revealed many recombination events that enabled initial delineation of a 1.25 Mb interval, which was flanked by the markers M54.19 and M55.45 (Fig. 4b, Table S2). All marker names are indicative of their genomic position: for example, M55.19 is located at 55.19 Mb on chromosome 4 of the reference genome BTx623 V3.1.1. To narrow the candidate region, two additional markers, M55.11 and M55.12, were developed (Fig. 4c; Table S2). Progenies of one of the key recombinant plants (R-39) showed true-to-type resistant response and all of them were homozygous for markers M55.11 and M55.12. In contrast, M55.09 was heterozygous in the susceptible plants and was thus excluded. Progenies segregated for the other key recombinant (R-51) although M55.12 was homozygote. Thus, M55.12 was also excluded from the *ARG3* genomic interval. Altogether, the recombination analyses supported placement of *ARG3* in a 30 kb genomic interval, flanked by M55.09 and M55.12 (Fig. 4).

**Fig. 4.**
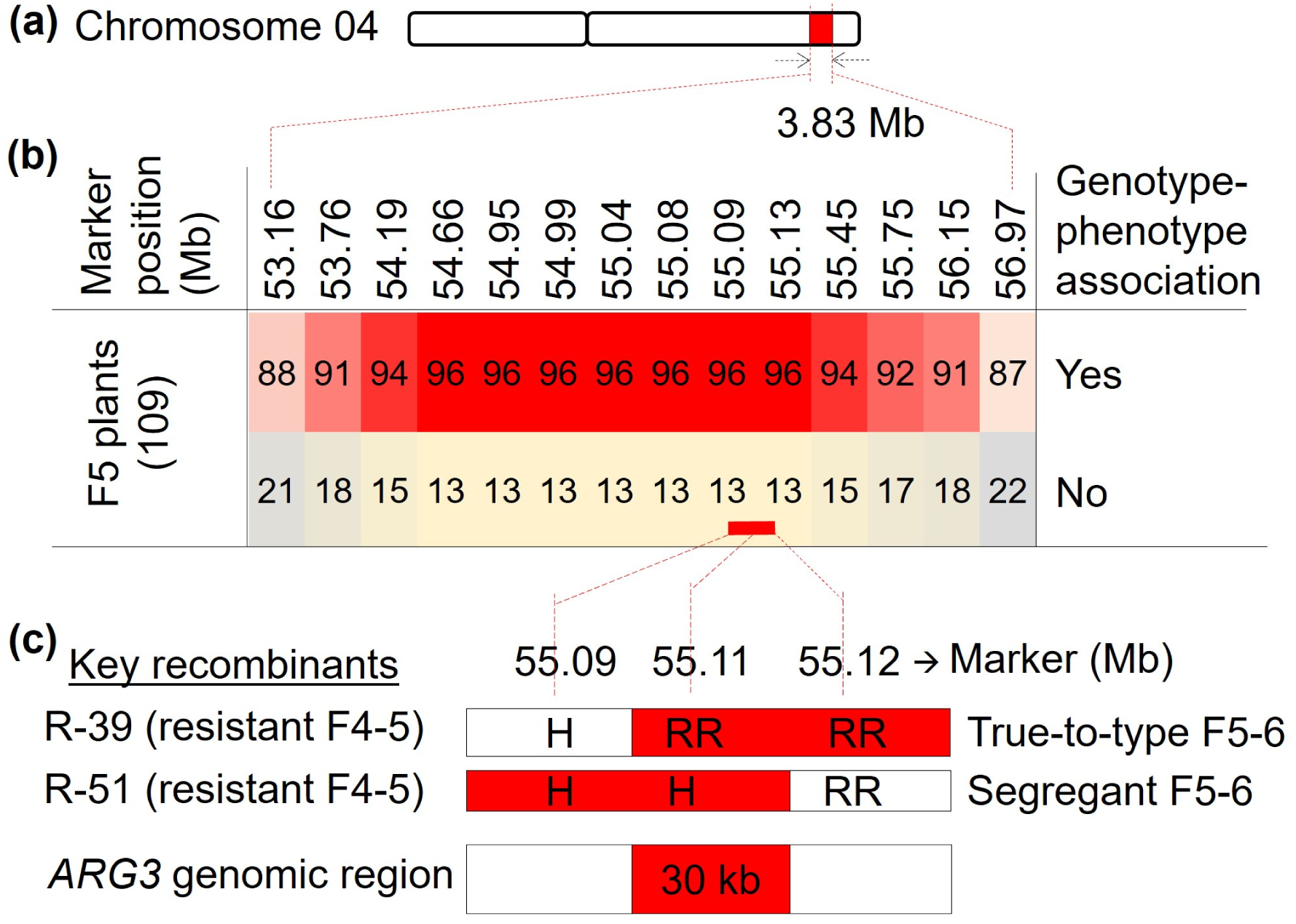
Fine map of the *ARG3* locus. (a) The *ARG3* locus maps to the distal end of chromosome 4. (b) Recombination analyses in 109 F4-5 (F4 derived F5 plants) across a 3.83 Mb genomic interval. Indel sequence markers are labeled by genomic position (53.16 indicates a marker M53.16 at 53.16 Mb). Detailed recombination data are provided in Table S2. (c) linkage analysis in two key recombinant resistant F4-5 plants. R-39 exhibited consistent resistance in the F5-6 generation, while R-51 segregated into resistant and susceptible individuals, enabling localization of the *ARG3* locus to a 30 kb interval. RR = homozygous resistant; H = heterozygous. The red shade marks the *ARG3* mapping intervals.

In addition to SNPs, there were structural differences in *ARG3* genomic region between parental lines used to generate the mapping population. Notably, both IS18760 and the resistant-bulk sample lacked a 12.4 kb repetitive sequence as identified using binary alignment map (BAM) files and genome assembly (Fig. 5a,b, Fig. S1a,b). Thus, the 30 kb recombination interval of the *ARG3* genomic region in BTx623 and TAM428 was only 17.7 kb in IS18760. The contigs carrying *ARG3* in TAM428 and the susceptible-bulk sample carried a portion of the 12.4 kb repetitive segment but not the non-repetitive sequence from the other side of the indel likely due to difficulty with *de novo* assembly of this region.

**Fig. 5.**
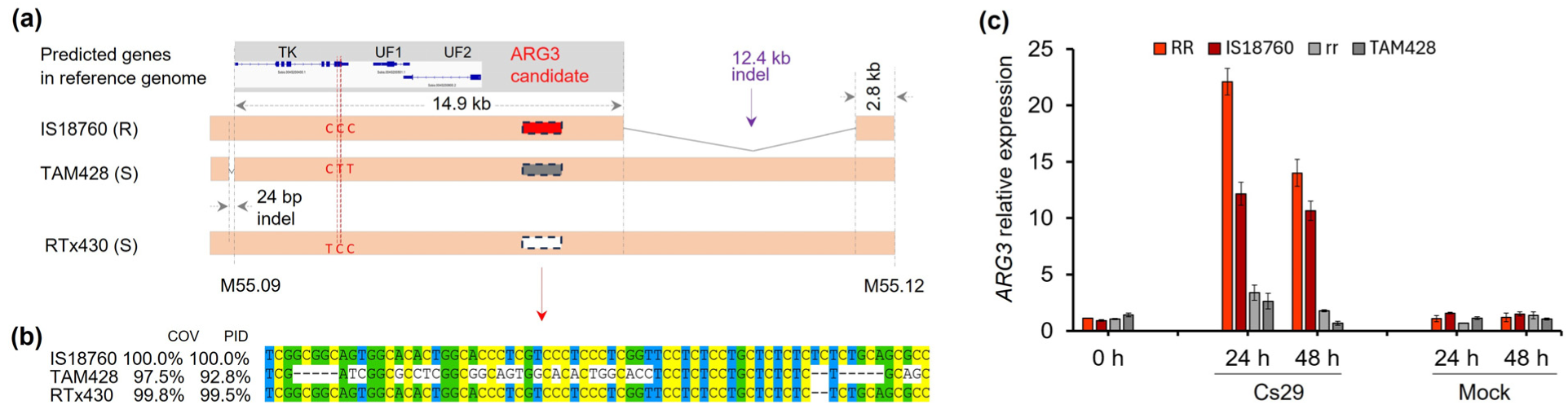
Polymorphisms in the *ARG3* genomic interval and expression of the *ARG3* candidate gene. (a) Genomic view of predicted genes in BTx623 sorghum reference genome v3.1.1: 17.7 kb (14.9 kb + 2.8 kb), in the resistant parental line IS18760 and 30 kb in the susceptible parental line TAM428. The region is flanked by markers M55.09 and M55.12. IS18760 and TAM428 differ by two SNPs outside of conserved region in TK (Tyrosine kinase gene). For TK, in contrast to TAM428, IS18760 and the susceptible line RTx430 vary with a single SNP in the coding sequence and share a 24 bp indel located at 1.3 kb upstream. Red, gray and white in ARG3 candidate region indicates the resistant genotype and two different susceptible genotypes. (b) A segment of aligned sequences of the expressed *ARG3* candidate gene in IS18760, TAM428, and RTx430. Coverage (COV) and percent identity (PID) compare sequences in the expressed 1,069 bp transcript. The complete alignment is presented in Supplementary File 2. (c) Relative gene expression (qRT-PCR) in parental lines and nearly isogenic lines. Leaf tissue samples were collected before inoculation (zero timepoint) and at 24- and 48-hours after Cs29 and mock inoculation.

### A rare allele of a protein non-coding *ARG3* gene confers anthracnose resistance

To define the *ARG3* candidate genes, the 30kb genomic region was examined for DNA sequence differences and functional domains of proteins in the mapping interval in the parental lines of the mapping populations. Binary alignment (BAM) files showed that the susceptible line BTx623, which was used as the reference genome, carried identical sequence in the region to that in the susceptible parental line TAM428, which facilitated the analysis of candidate genes (Fig. 5, Fig. S1). In these backgrounds, the region harbors three predicted genes: Sobic.004G200400, which encodes a tyrosine kinase domain protein, and two others (Sobic.004G200501 and Sobic.004G200600), which carry domains of unknown function.

Comparison between the two parental alleles of Sobic.004G200400 revealed that they differ in two SNPs that changed proline to serine and threonine to methionine in the susceptible parent relative to the resistant line, and the third SNP encodes for a synonymous amino acid (Fig. 5a, Fig. S2a,b). Both nonsynonymous SNPs are outside the tyrosine kinase domain, and neither of these changes are to the conserved residues (Fig. S2). In addition, the parental lines differ in a 24 bp indel 1.3 kb upstream, which is absent in the susceptible parent. However, another susceptible line, RTx430, shared both the nonsynonymous amino acids and the upstream indel sequence with the resistant parental line IS18760 (Fig. 5a). The gene carried a single SNP between RTx430 and IS8760 (Supplementary File 1). The gene was downregulated in response to the pathogen inoculation although it showed higher basal transcript level in the resistant parental line compared to the susceptible parental line, and the plantNexus database, built using hundreds of sorghum variants, shows a high relative transcript level throughout the plant. Collectively, these data suggest that this gene is not a likely candidate *ARG3* gene underlying the pathogen phenotypes observed.

The other two candidate genes (Sobic.004G200501 and Sobic.004G200600) were identical in the parental lines as revealed by aligned genomic raw-read data and genome assembly (Fig. 5a, Fig. S1a). In addition, transcripts of these genes were not detected in healthy and pathogen inoculated leaf tissue samples of the parental lines. In the PlantNexus transcriptome database, the Sobic.004G200501 transcript was detected in the root and reproductive tissue samples but not in the leaf, where the host-pathogen interaction occurs (Fig. S3a). Similarly, Sobic.004G200600 showed no transcript in the vegetative tissue (Fig S3b). Thus, our analysis supported that these two predicted genes are unlikely *ARG3* candidates.

The remaining *ARG3* genomic region lacks any putative (annotated) gene but a signature of aligned transcript raw-reads in the sorghum reference genomes and showed unusually high polymorphism between the parental lines IS18760 and TAM428 (Fig. S1, Fig. S4). Using RT-PCR, we found a 1,069 bp RNA sequence in the resistant parental line that was identical to the genomic sequence and showed several indel sequences and SNPs between the parental lines (Fig. S4). Another susceptible line, RTx430, carried distinct polymorphisms in this expressed 1,069 bp sequence (Fig. 5b, Supplementary File 2). This transcript was strongly induced 24 h and 48 h after inoculation in the resistant nearly isogenic lines and the resistant parental line, and to a much lower extent in the susceptible line (Fig. 5c). The extensive sequence divergence between the parental lines, as well as early induced expression in the resistant lines is consistent with a resistance response and suggested that the expressed gene lacking annotation was a strong candidate for the *ARG3* gene.

In the resistant parental line (IS18760), the *ARG3* sequence (1,084 bp) contains no open reading frame longer than 243 bp (80 codons, reverse strand) across either strand or any of the six reading frames which is shorter than would be expected for a functional protein coding gene. Sequence comparison between the resistant (IS18760) and susceptible (TAM428) alleles revealed that divergence at this locus is overwhelmingly indel-driven: 15 independent indel events, ranging from 1 to 17 bp, compared to only 5 single-nucleotide substitutions across the aligned region. This pattern is inconsistent with purifying selection acting on a protein-coding gene, where indels are strongly disfavored due to their frameshifting consequences and point substitutions typically dominate, thus is consistent with a locus evolving free of coding constraint. Together with the absence of any substantial ORF and the lack of recognizable protein domains or cross-kingdom homology for the candidate short ORFs, these results support *ARG3* as a non-coding RNA gene rather than a protein-coding gene (Fig. S4). In addition, based on analysis using the Coding Potential Calculator 2 (https://cpc2.gao-lab.org/index.php), *ARG3* was classified as a non-coding sequence, with a coding probability of only 0.14, indicating ∼ 86% likelihood of being non-coding. None of 111 *sorghum* pangenomes carried sequences identical to that in the resistant parental line, suggesting that it is a rare allele that confers resistance (Fig. 6a).

**Fig. 6.**
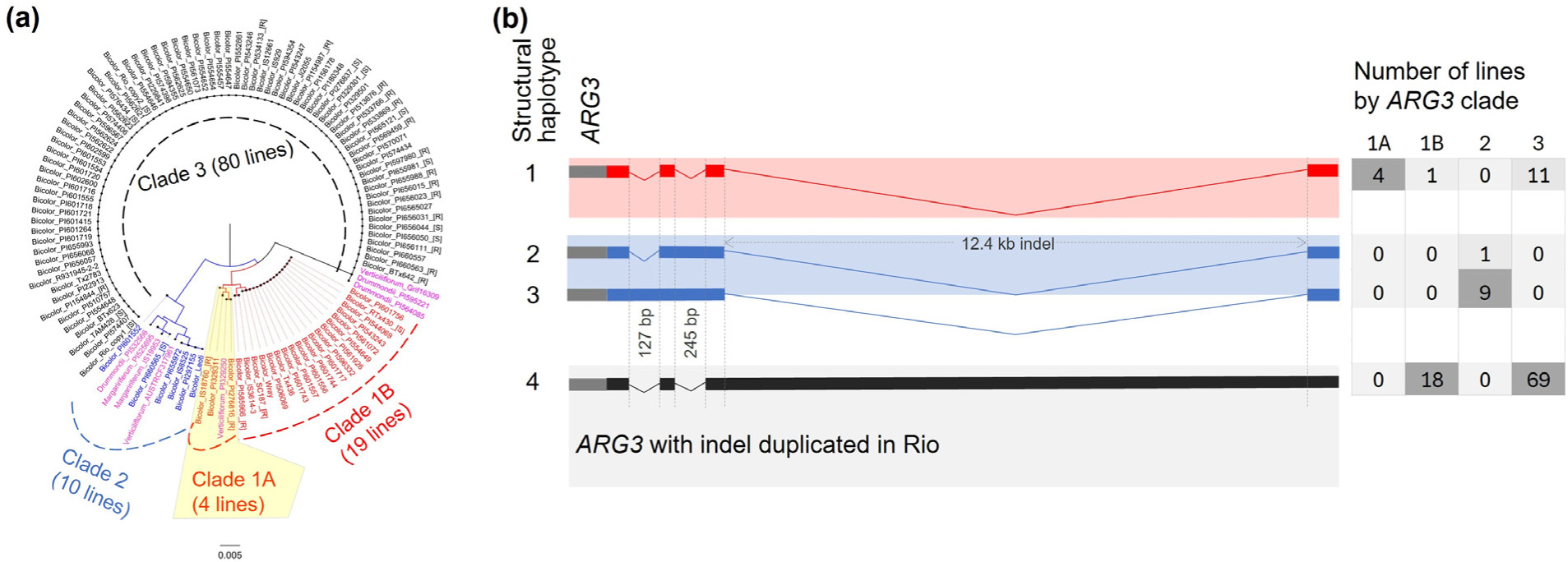
Phylogenetic analyses of the *ARG3* nucleotide sequence in the sorghum pangenome and the structure of the ARG3 locus (indel sequences). (a). Phylogenetic analyses the *ARG3* sequence in 111 sorghum pangenome lines with the parental lines IS18760 and TAM428. In accession labels, the prefixes “Bicolor” and others such as “*Drummondii*” for wild sorghum indicate subspecies, and the suffix [R] for resistant and [S] for susceptible show response of 28 lines to Cs29 in addition to the parental lines of the mapping population. The rare variant in the resistant parental line, IS18760, was clustered in clade 1A, and the susceptible parental line TAM428 was clustered in clade 3. (b) Distribution of the selected three indel sequences in clades of the *ARG3* sequence. *ARG3* sequence with the 12.4 kb sequence shows duplication in Rio reference genome.

### Evolutionary insights concerning the *ARG3* locus

*ARG3* sequence variation and phylogeny were examined in 111 *sorghum* pangenome lines (Fig. 6a). These include 80 genomes available through *sorghumbase.gov* and additional 31 genomes extracted from the 37-pangenome in *phytozome-next.jgi.doe.gov*, most of which we evaluated for responses to *C. sublineola* strain Cs29 (Fig. 6a). The allele of *ARG3* in the susceptible parental line TAM428 was carried by 68% of the pangenome lines but no line carried sequences identical to that in the resistant parental line IS18760. Twenty lines carried >99.4% nucleotide identity to the allele in this parent with 99.8% coverage, and in the 88 lines identity falls to <94%. Sequence alignment of 18 representative lines and *Sorghum bicolor* subspecies are presented in Supplementary File 2. Several indel sequences of a few-to-many base pairs were found in the expressed 1,069 bp *ARG3* sequence (Fig. S4, Supplementary File 2), which was a major contribution to the overall nucleotide similarity (Fig. 6a). *ARG3* nucleotide sequences clustered into three main clades with IS18760 in clade 1 and TAM428 in clade 3 (Fig. 6a). Wild sorghum *(Sb. subsp. drummondii*, *verticiliflorum,* and *marganiriferum*) was represented in all clades and was relatively enriched in clade 2, which is more closely related to clade 1 than to clade 3 (Fig. 6a).

Several indel sequences were found in the potential cis-regulatory sequences as well. Three indel sequences (127 bp, 245 bp and 12.4 kb) in the pangenome were examined for relatedness to ARG3-mediated resistance response (Fig. 6b). The 127 bp and 245 bp indel sequences were 1 kb upstream of the expressed 1,069 bp *ARG3* sequence, and their presence was exclusive to clade 2 and was always found in the absence of the 12.4 kb sequence. Both the 127 bp and 245 bp indel sequences were absent in the parental lines of the *ARG3* mapping population as well as in the entire clade 1 and clade 3 to which both parental lines belong. Thus, we conclude that they have no role in gene expression differences between the parental lines, although they may have disrupted *ARG3* function in clade 2 lines, which are enriched in sequences from wild sorghum accessions.

All four lines of clade 1A (three *Sb. subsp. bicolor* accessions, including IS18760 and one *Sb. subsp. verticiliflorum*) lack the 12.4 kb indel sequence. These four lines as well as the pathogen strain Cs29 originated in Ethiopia, consistent with co-evolution of resistance with the presence of Cs29 which is avirulent on IS18760. All but one of 19 lines of clade 1B, including the susceptible RTx430, carried the 12.4 kb insertion sequence, which was absent in all four lines of clade 1A (Fig. 6a), which supports association of the absence of this insertion with ARG3-mediated resistance. In addition, presence of this indel sequence in 86% (69 of 80) lines of clade 3 carrying the susceptible parental line TAM428 supports the hypothesis that the insertion is related to the lack of resistance.

It is notable from analysis of the pangenome, in addition to *ARG3*, that other loci that were not considered in this study also likely confer resistance to Cs29, which is evident by differences in host responses of lines carrying identical *ARG3* sequences (Fig. 6a); 23 lines in clade 3 carry identical *ARG3* sequences, but 15 were resistant and eight were susceptible, suggesting that in these eight lines, *ARG3* was not sufficient to confer resistance. Similarly, in clade 1B, the susceptible line RTx430 and the resistant line SC187 carry identical *ARG3* sequence suggesting the presence of other major loci in other backgrounds that mediate resistance to Cs29.

### The *ARG3* non-coding RNA gene is required for resistance to *Colletotrichum sublineola*

To validate that *ARG3* non-coding RNA gene is required for resistance to *C. sublineola* and is the causal gene for resistance at the *ARG3* locus, we used virus-induced gene silencing (VIGS) to suppress *ARG3* gene expression in the resistant IS18760 background and tested for responses to the pathogen. VIGS was implemented as described previously ((Singh *et al*., 2018; Mei *et al*., 2019) and in the methods section) using a fragment of the *ARG3* gene. qRT-PCR confirmed a significant reduction in *ARG3* transcripts in the silenced lines (IS18760_1, IS18760_2, IS18760_3, and IS18760_4) relative to IS18760 plants inoculated with the empty vector (IS18760_EV1, IS18760_EV2, and IS18760_EV3) (Fig. 7a). Inoculation of the VIGS-treated IS18760 plants with *C. sublineola* caused enhanced disease symptoms and increased fungal accumulation, as measured by qPCR of fungal genomic DNA amplified using *C. sublineola*-specific primers, in the *ARG3*-silenced lines compared with the empty-vector control (Fig. 7b, c). The loss of resistance in IS18760 after silencing, and the data in the preceding sections, collectively suggest that *ARG3* is a rare allele of a long non-coding RNA gene that is required for resistance to *C. sublineola*.

**Figure 7.**
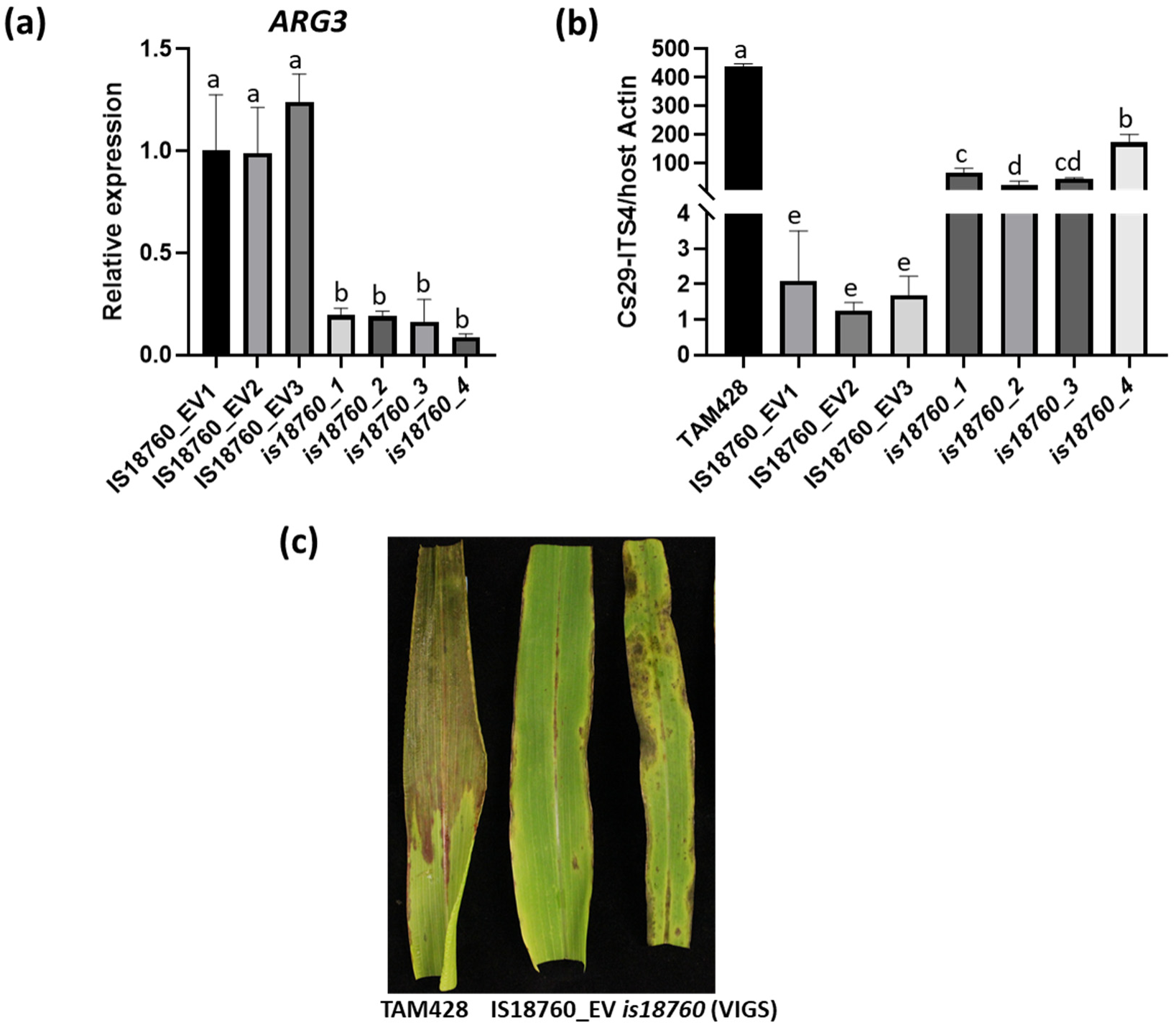
The *ARG3* non-coding RNA gene is required for resistance to *Colletotrichum sublineola.* a) Gene expression, (b) Disease symptom, and c) Fungal growth after silencing of *ARG3* non-coding RNA gene in the background of the *Colletotrichum sublineola* resistant sorghum line IS18760. In (a) Expression levels of *ARG3* were analyzed by qRT-PCR in three sorghum IS18760 plants rub inoculated with an FoMV empty vector (IS18760-EV1, IS18760-EV2 and IS18760-EV3) and four plants rub inoculated with the ARG3-VIGS vector (*is18760-1, is18760-2, is18760-3* and *is18760-4*) at five days after *C. sublineola* inoculation. The data obtained was normalized by the comparative cycle threshold method with sorghum *Actin* as the internal control and presented as relative expression. Error bars show standard deviation (*n* = 3 biological replicates each performed on RNA extracted from individual whole plants); statistical differences were determined using a one-way ANOVA followed by a Tukey test. Experiments were conducted two times with similar results. In (b) disease response symptoms are at 7-day post inoculation with *C. sublineola* showing TAM428 (susceptible control line), IS18760-EV and *is18760*. In (c) qPCR quantification of fungal growth. Genotypes are the susceptible sorghum background TAM428, control treatment (FoMV empty vector in IS18760: IS18760-EV1, IS18760-EV2 and IS18760-EV3), after silencing treatment with the *ARG3* non-coding RNA gene in the IS18760 background (*is18760_1*, *is18760_2*, *is18760_3* and *is18760_4*). Fungal growth in infected leaves was determined by qRT-PCR amplification of the *C. sublineola ITS* region (Cs-ITS). Relative DNA abundances were calculated using sorghum *Actin* (*Sb Act*) as the reference gene. Error bars show standard deviation (*n* = 3 biological replicates each performed on genomic DNA extracted from individual plants). Statistical differences were determined using a one-way ANOVA followed by a Tukey test.

### Synteny and comparative analysis of *ARG3* non-genic regulation

*ARG3* showed no significant homology to annotated genes or mapped transcripts in the reference genomes of related species (Fig. S5). However, many genes flanking *ARG3* are syntenic in rice and several other grasses (Fig. 8a). A database of various histone modification as well as DNA methylation (https://epigenome.genetics.uga.edu/PlantEpigenome/index.html) was used to examine clusters of conserved non-coding sequences, focusing on those with the highest levels of conservation, confirmed using the CoGe comparative genomics tool (https://genomevolution.org/coge/) (Lyons *et al*., 2008; Lu *et al*., 2019; Ricci *et al*., 2019). A notable feature of the *ARG3* locus is that it is a region rich in conserved non-coding sequences. When compared to syntenic regions in other grass species, including *Oryza sativa*, *Setaria viridis*, *Zea mays*, and *Brachypodium distachyon*, there are patterns of conservation consistent with conserved function, albeit not conserved coding-potential (Fig. 8b). Notably, there are also conserved chromatin marks (Fig. 8b). Remarkably, we found that in each case, the non-coding region that was best conserved was depleted with respect to DNA methylation and enriched in H3K27me3, and H2A.Z, consistent with facultative repression of gene expression, and specifically with environmentally responsive genes (refs) (Fig. 8b). These data suggest that the chromatin features associated with *ARG3* have been conserved since at least the split between rice and sorghum.

**Fig. 8.**
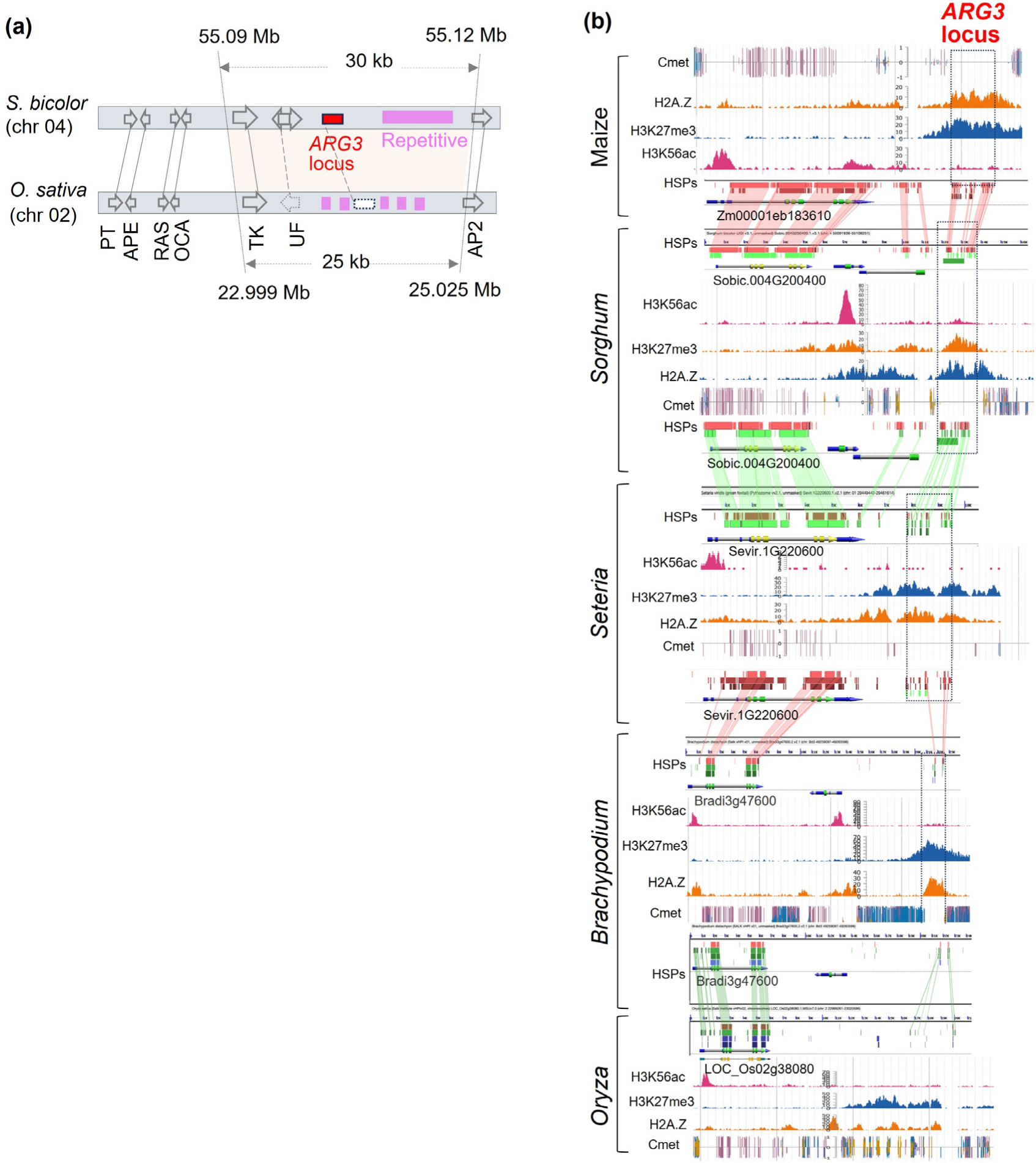
Synteny in the ARG3 genomic region, and histone and DNA modifications. (a) Syntenic genes between sorghum and rice: Phosphate transporter (PT), anaphase promoting element (APE), RAS suppressor (RAS), Ovarian carcinoma antigen related (OCA)/BRCA1, methyltransferase (MT), Tyrosine kinase (TK), and AP2 domain (AP). Solid line connecting the panels shows homologous genes, and dash line connects weak sequence similarity where putative gene was not found in one or both species. Purple color shows repetitive sequences in ARG3 locus. Light brown shade spans the 30 kb ARG3 genomic region. (b) Comparative genomics and histone and DNA modification in rice, Brachypodium, Setaria, sorghum, and maize. High score pairs (HSP) sequences indicate regions of sequence similarity. Levels of specific histone modifications for this region in each genome are as indicated in orange, blue, and purple.

### Additional anthracnose resistance QTL identified using the TAM428 x IS18760 mapping population

F1 plants showed the resistance response with the extensive cell death phenotype, and in nearly isogenic lines, *ARG3* is a major contributor to anthracnose resistance and is associated by extensive cell death (Fig 1c, Fig. 3a). The association of *ARG3*-phenotype that restricted pathogen growth (Fig. 3a) with the parental alleles indicated that the resistance was inherited as a dominant allele (Table S2). Whereas the extensive cell death in F1 plants (Fig. 1c) suggest that the phenotype is dominant over that in the IS1876-type resistance response, which lacks the extensive cell death (Fig. 1a,b).

In contrast, the cell death phenotype in IS18760 was highly restricted (Fig. 1a,b), suggesting the presence of other loci that contribute to the resistance phenotype. Although the statistical power was low compared to that of *ARG3*, BSA-Seq analysis showed additional QTL, including one on the first arm of chromosome 04 and a second one on distal end of chromosome 10 that contributed to the resistance in IS18760 (Fig. 2b, Fig. S6). The sliding-window based ΔSNP-index is a powerful QTL analytical method because it provides estimates of SNPs in a wide (2 Mb or 4 Mb) genomic intervals. Thus, QTL with approximately 0.05 p-value is less likely to occur by chance. The QTL on chromosome 10 harbors a BTB/POZ domain predicted gene (Sobic.010G227100) that encodes a BTB/POZ domain protein family. Its ortholog in rice LOC_Os06g46240 confers resistance to bacterial leaf blight disease (Fatimah *et al*., 2018), and its homologs were implicated in signaling roles in response to pathogens (Zhang *et al*., 2019; Zhou *et al*., 2022; Mandal *et al*., 2023). Genomic data (BAM files) of IS18760 and TAM428, and their progenies that were bulk sampled showed a 32 bp indel sequence 100 bp upstream of the gene IS18760. that may influence gene expression and complement *ARG3* for the restricted cell death resistance.

## Discussion

The sorghum line IS18760 exhibits broad-spectrum resistance to the highly virulent *Colletotrichum sublineola* strain Cs29 and to many other isolates that cause anthracnose disease in sorghum (Mewa *et al*., 2023; Mekonen *et al*., 2024). To date, the genes underlying this resistance have not been identified. To investigate the genetic basis of Cs29 resistance in IS18760, we developed a biparental mapping population derived from a cross with the highly susceptible line TAM428. BSA-seq resulted in multiple loci contributing the resistance response, with fine mapping pinpointing a major locus on chromosome 4, which we designated, *ANTHRACNOSE RESISTANCE GENE3 (ARG3)*. ARG3-mediated resistance is associated with extensive host cell death that restricts pathogen proliferation. The *ARG3* locus was fine mapped to a 30 kb genomic region carrying three predicted genes and one unannotated gene. The three predicted genes did not exhibit DNA polymorphisms or carried polymorphisms that were inconsistent between the resistant and susceptible lines. However, the unannotated non-coding RNA gene exhibited significant sequence polymorphism between the resistant and susceptible lines and induced with a higher transcript level in the resistant line after pathogen inoculation. This pattern of gene expression in the single *ARG3* candidate gene is consistent with resistance response against the pathogen (Wharton & Julian, 1996; Mewa *et al*., 2023). Sequence analyses in *sorghum* pangenome identified other recessive alleles of *ARG3* in susceptible lines, while the resistant allele appeared to be rare. The identification of the *ARG3* locus is consistent with previous reports that localized anthracnose resistance to the same genomic region in IS18760. A 2 Mb region on chromosome 4 (spanning 65-54 Mb), which includes the *ARG3* locus (Fig. S7), was previously reported to harbor an anthracnose resistance QTL (Patil *et al*., 2017; Cuevas *et al*., 2021).

Multiple lines of evidence indicate that ARG3 functions as a non-coding RNA rather than a protein-coding gene. Sequence analysis revealed no significant open reading frame, and divergence between resistant and susceptible alleles consists mainly of indels rather than single-nucleotide substitutions,a pattern inconsistent with purifying selection on a coding sequence, where indels are strongly disfavored due to frameshifting. Most directly, VIGS silencing of the ARG3 transcript rendered the resistant line susceptible to *C. sublineola*. Since VIGS acts through sequence specific RNA degradation independent of translation or reading frame, this result demonstrates that the *ARG3* RNA itself but not a translated product is required for resistance. Together, the absence of an ORF, the indel-dominated divergence pattern, and the VIGS phenotype provide direct evidence that RNA-level function underlies *ARG3*-mediated resistance in the sorghum line IS18760.

The *ARG3* allele in IS18760 is a rare variant as we did not find identical sequence in 111 diverse *Sorghum* pangenome. The Ethiopian core germplasm collection showed an abundance of rare variant SNPs in the range of 46-60% (Cuevas *et al*., 2017; Girma *et al*., 2019). IS18760, the a wild sorghum accession carrying one of the closest *ARG3* sequences to it as well as the pathogen strain Cs29 originated in the *S. bicolor* domestication region in East Africa including Ethiopia and Sudan, which entail the possibilities of wild to cultivated gene introgression and *ARG3* to Cs29 evolutionary link (Prom *et al*., 2012; Venkateswaran, 2019). Cultivated sorghum has undergone gene exchange with the weedy types belonging to wild sorghum subspecies (de Wet, 1978), which is consistent with our observation that one of the closest known DNA sequence similarity to the resistant allele was in the *Sb. subsp. verticiliflorum* line named PI329250. Indeed, *ARG3* sequence variation was not discernibly different between the cultivated and wild lineages, which is consistent with a possible gene exchange between them.

Structural variation associated with repetitive sequences drives the evolution of plant genomes and often influences stress related genes (Pinosio *et al*., 2016; Fuentes *et al*., 2019). *ARG3* and the flanking sequence showed high structural variation, including a 12.4 kb indel of repetitive sequence, which was absent in the resistant line and whose absence showed evolutionary relatedness to the resistance. ARG3-mediated resistance might be disrupted due to insertion of this repetitive sequence. This degree of within-species sequence variation is unusual for a protein coding gene (Mattick *et al*., 2023). The unusually high sequence polymorphism, lack of a conserved open reading frame and absence of homology to known proteins suggests that *ARG3* is a non-coding RNA gene that confers selective advantage and have been maintained over time through regulation of other genes (Ruiz-Orera *et al*., 2020). On the other hand, many non-coding RNAs encode peptides (Rion & Rüegg, 2017), and we have not ruled out that possibility for *ARG3*. The lack of *ARG3* gene annotation in sorghum and absence of orthologous genes in related organisms restrict possibilities for further studies based on comparative genomics insights. However, the presence of a clear set of syntenic conserved non-coding sequences suggests a conservation of DNA-mediated changes. It would be worth examining gene expression of this region in other species following pathogen exposure using raw sequence data rather than annotated gene models to see if this syntenic region in those species also encode non-coding RNAs.

Cs29 appears to be virulent to a large proportion of the sorghum germplasm, including the broad-spectrum anthracnose resistant sorghum line SC748-5 (Prom *et al*., 2012; Mewa *et al*., 2023). Nevertheless, IS18760 showed resistance to Cs29, and all tested 25 Cs isolates collected from various sorghum producing agroecologies in Ethiopia (Mekonen *et al*., 2024). It is of interest that both Cs29 and resistance in IS18760 to this strain is rooted in the western Ethiopia and eastern Sudan ecological belt wherein the species was domesticated (Morris *et al*., 2013). In contrast, IS18760 is susceptible to Cs strains that were collected in the USA (Mewa *et al*., 2023), which makes *ARG3* of interest with respect to the evolution of host resistance and pathogen virulence.

The discovery that *ARG3* is a non-coding RNA gene opens new possibilities for understanding the molecular basis of fungal resistance in sorghum. Although the mechanism of ARG3-mediated resistance remains to be elucidated, our findings suggest possible mechanisms based on known functions of non-coding RNAs in plant immunity (Song *et al*., 2021). ARG3-mediated resistance to sorghum anthracnose disease was inherited as a dominant allele, and the functional allele is induced by infection and confers fungal resistance. This suggests that ARG3 may act as a regulatory RNA that modulates expression of defense-related loci either in *cis* or *trans*, potentially by recruiting chromatin-modifying complexes or interacting with transcriptional regulators to promote an active chromatin state (Han & Chang, 2015). It is also possible that ARG3 suppresses negative regulators of defense, serves as a molecular scaffold or RNA decoy, binding and sequestering immune-suppressive microRNAs to de-repress defense transcripts involved in the hypersensitive response and reactive oxygen species (ROS) production (Franco-Zorrilla *et al*., 2007). The observed phenotype of enhanced cell death due to the *ARG3* locus supports a role of the gene in amplifying downstream cell death pathways. Further experiments, including identification of regulators of induced *ARG3* gene expression, proteins that interact with the *ARG3* transcript, and the target loci that *ARG3* regulates are essential to clarify the mechanism by which ARG3 contributes to durable resistance against *C. sublineola* but are beyond the scope of this work.

The current study identified and characterized a previously unannotated non-coding sorghum anthracnose candidate resistance gene through high-resolution genetic mapping, pangenome data analysis, gene expression, and in silico data. Given the attributes of *ARG3* described in the current study, ARG3***-***mediated host resistance might uniquely contribute to disease management and offer novel insights into host resistance mechanisms in crop plants. Further investigation for new resistance alleles as well as understanding the distribution of Cs pathotypes against ARG3-mediated plant immunity might enhance usability of the gene for resistance breeding.

## Acknowledgements

We thank former and current members of the Mengiste lab, including undergraduate students, for their assistance in maintaining plant materials. We also thank the Purdue Herbaria for access to microscopy facilities used to capture images of trypan blue–stained tissues. D.L was supported by Hatch funds.

## Author Contributions

DBM conducted most of the experiments including identification of the resistant parental line, population development, data collection, analysis, and writing the paper. AA contributed to the genetic analysis and gene expression analysis of *ARG3* and edited the manuscript. AG performed gene expression analyses, conducted all the steps of the virus induced gene silencing from constructs to diseases assays, and wrote and edited the manuscript. PO conducted trypan blue staining and repeated the phenotypic assays. CJL investigated histone methylation. GG contributed to the genetic analyses of the *ARG3* gene. ES contributed to *ARG3* gene expression and cDNA. MG contributed to genomic data analysis. DL conducted comparative analysis of *ARG3* histone modification and was involved in writing and editing the manuscript. TM directed the project, generated experimental ideas and wrote and edited paper.

## Data Availability

Whole-genome resequencing data, assembled contigs, and *ARG3* sequence files are available upon request or through public repositories associated with this study. Additional datasets supporting the findings of this work, including marker sequences, alignment files, and pangenome comparisons, are provided in the Supporting Information.

## Supporting Information

**Table S1.** BSA-Seq statistics in the *ARG3* genomic region.

**Table S2.** Marker genotypes of the recombination analysis in the *ARG3* QTL region.

**Table S3.** Primers used for qPCR of pathogen growth, recombination based fine mapping, gene amplification, and qRT-PCR

## Supplemental Figures

**Fig. S1.**
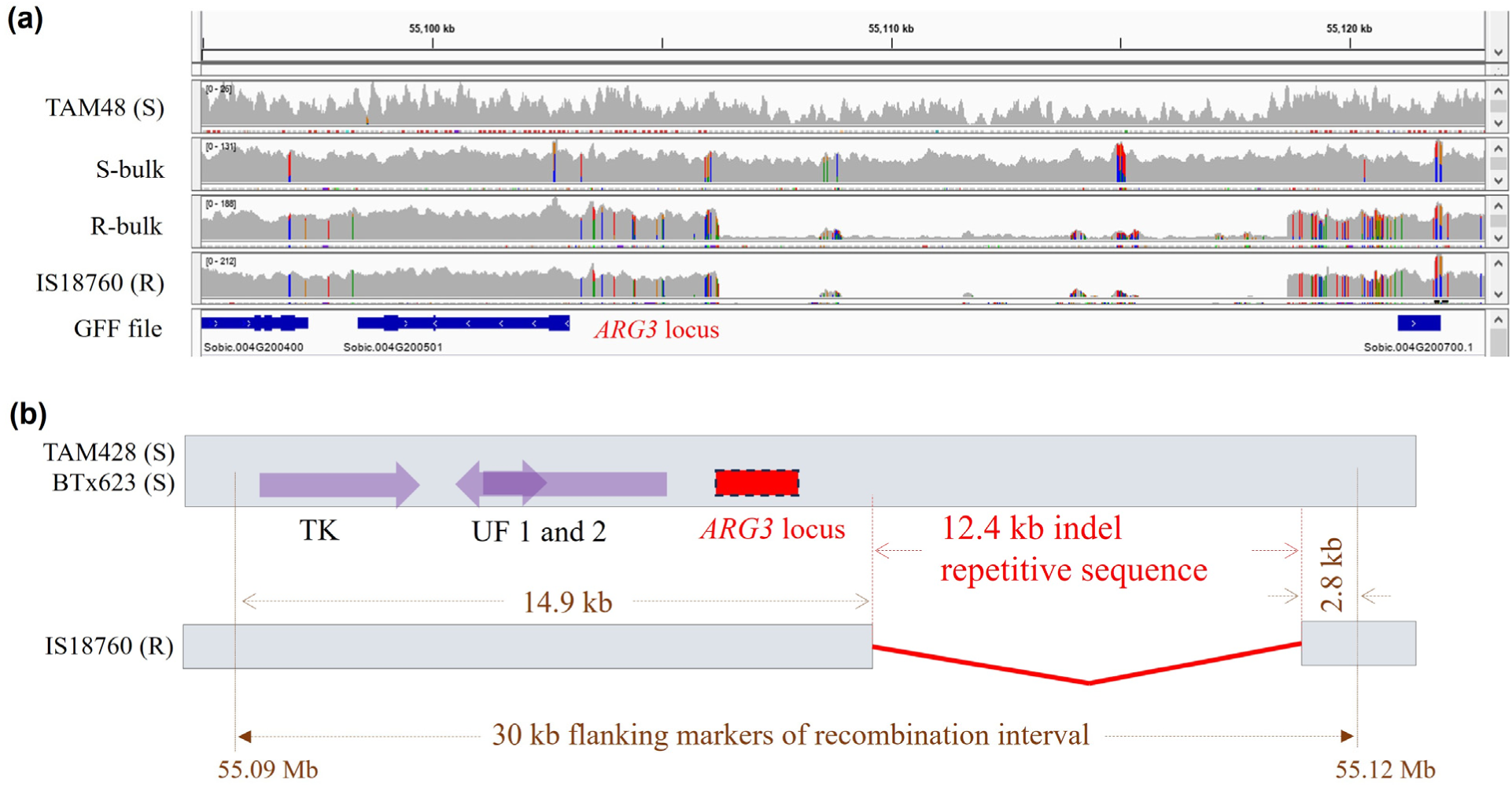
Variation in the 12.4 kb indel in the 30 kb *ARG3* genomic region. (a) Snippet of binary alignment (BAM) files in the parental lines and bulk samples in integrated genome viewer (IGV). The indel is absent in samples of the resistant genotypes. S-bulk = susceptible progenies bulk sample used in BSA-Seq. R-bulk = resistant progenies bulk sample used for BSA-Seq. The bottom panel is the annotation file of the reference genome. (b) A diagram showing the 12.4 kb indel in de novo assembled contigs.

**Fig. S2.**
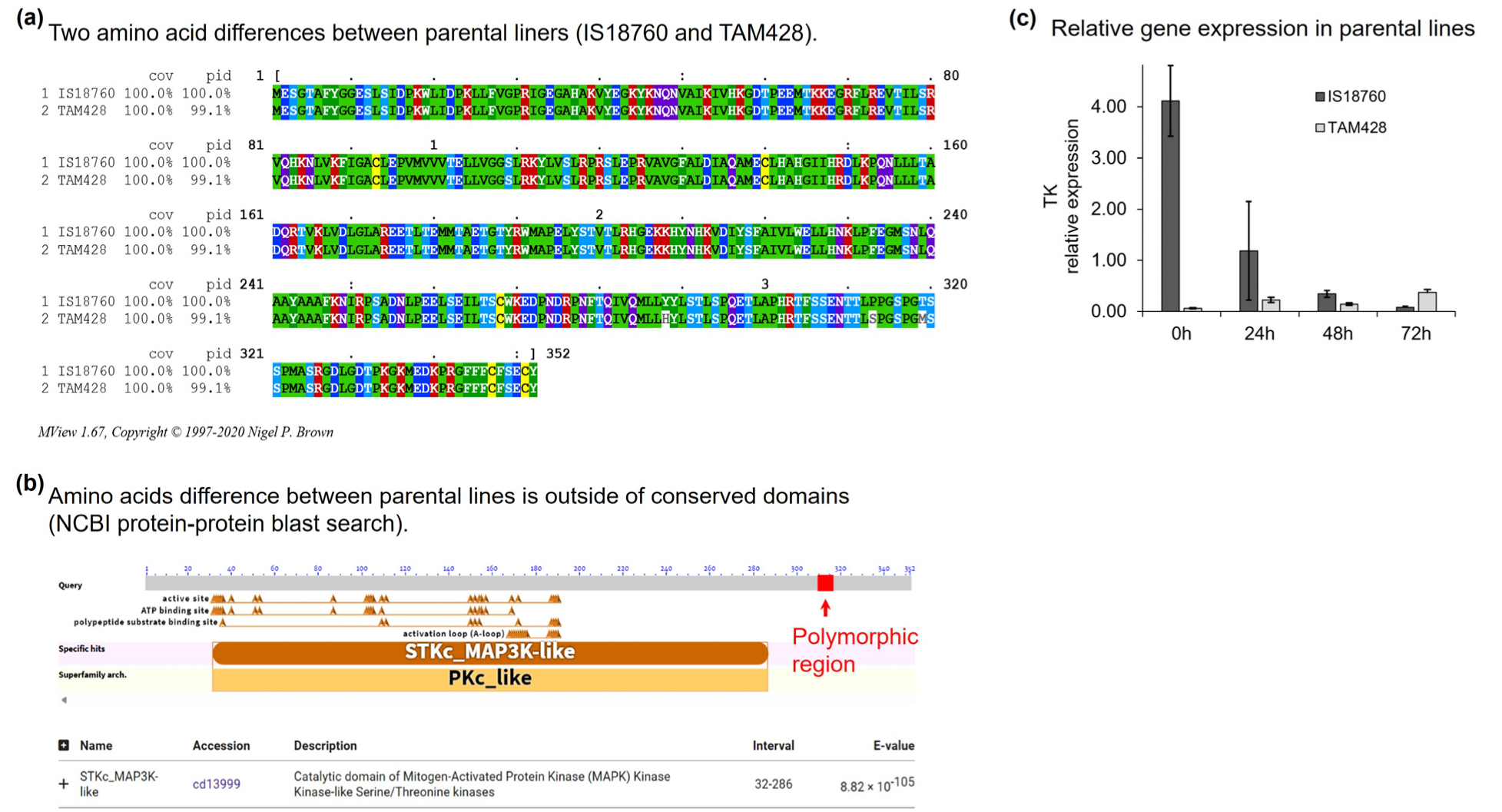
Polymorphism and gene expression in the non-candidate tyrosine kinase (TK) domain gene, which is found in the *ARG3* genomic map interval. (a) Polymorphism between parental lines. (b) All polymorphisms between the parental lines are outside the conserved domain. (c) The gene showed downregulation with pathogen (Cs29) inoculation.

**Fig. S3.**
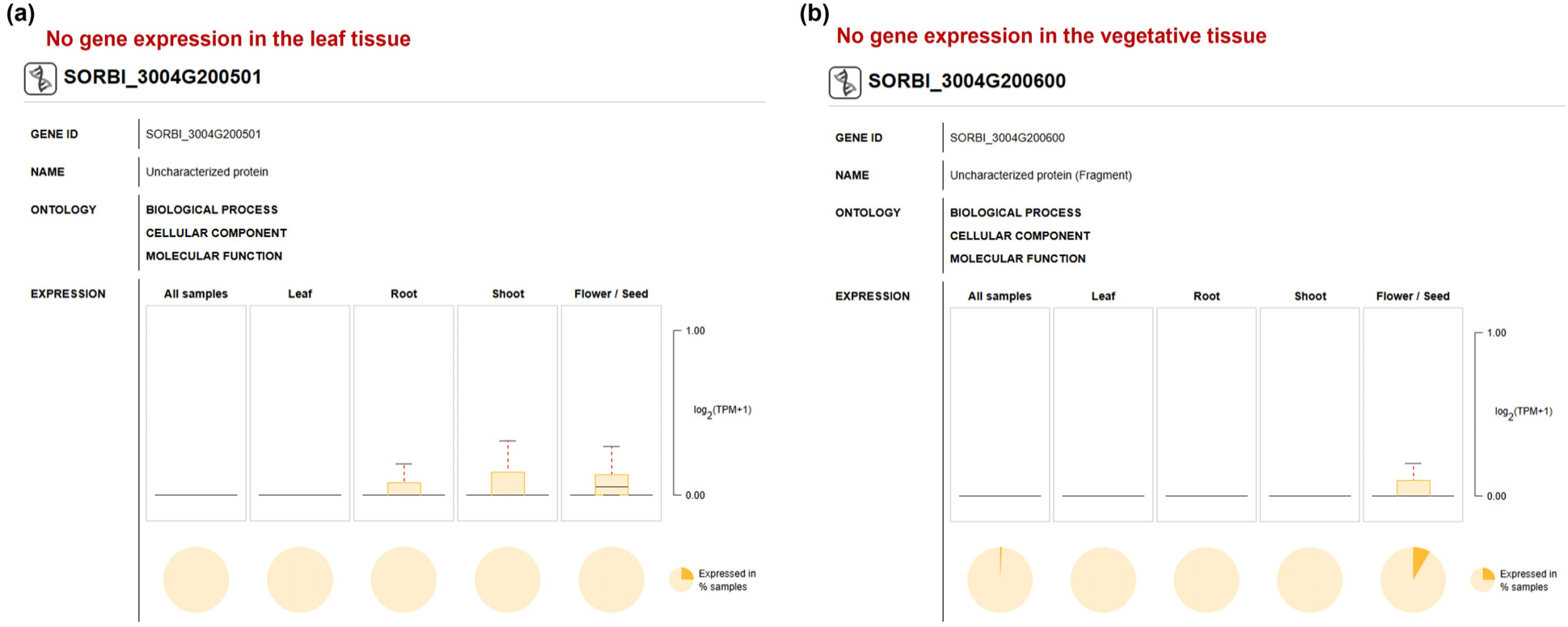
Expression profile of the two putative genes of unknown function UF1 (a) and UF2 (b) that reside in the 30 kb *ARG3* map interval from pan-transcriptome data in Plant Nexus database.

**Fig. S4.**
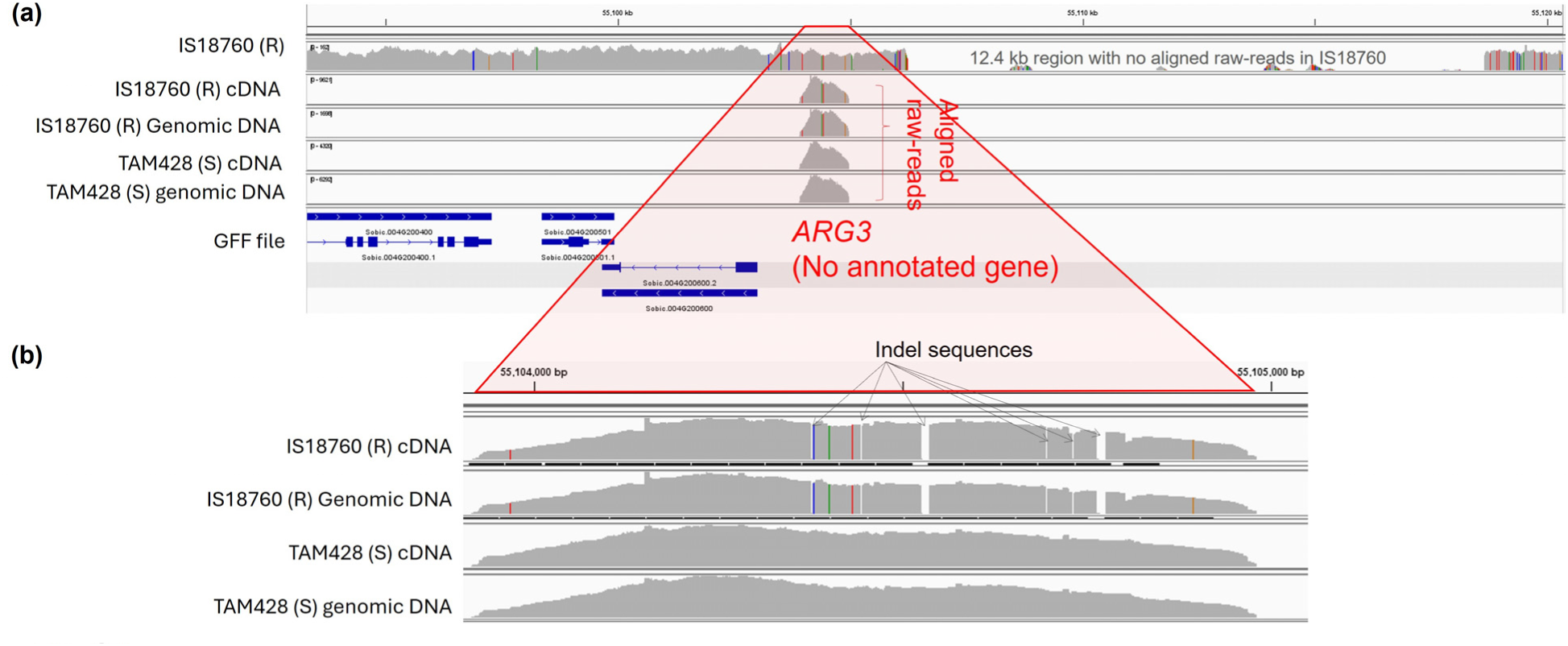
*ARG3* raw-reads aligned to the reference genome. (a) Top to bottom: snippet of BAM files of IS18760 re-sequence raw-reads, and Wide-Seq amplified sequences of IS18760 cDNA, IS18760 genomic DNA, TAM428 cDNA, TAM428 genomic DNA. Red, blue, orange and green vertical lines in the BAM files show nucleotides that are different from the reference genome to which the alignment was built on. The bottom panel shows putative genes in the reference genome and *ARG3* has no annotation. (b) Expanded view of the aligned *ARG3* raw-reads showing five indels.

**Fig. S5.**
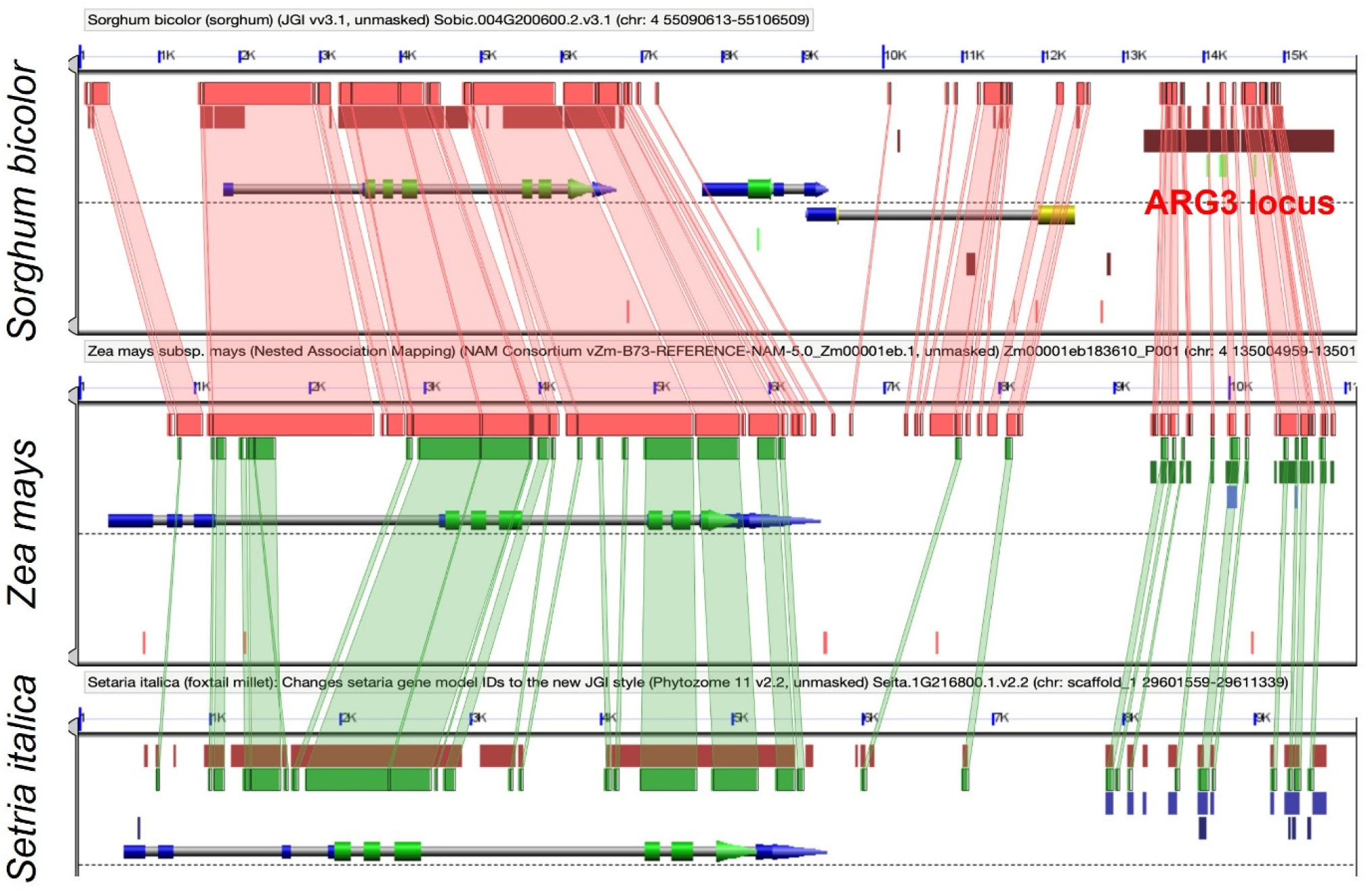
Homology of *ARG3* genomic interval with other grasses. Close up view of the homologous regions in the 30 kb region presented in Fig. 6. https://genomevolution.org/r/1qbkg

**Fig. S6.**
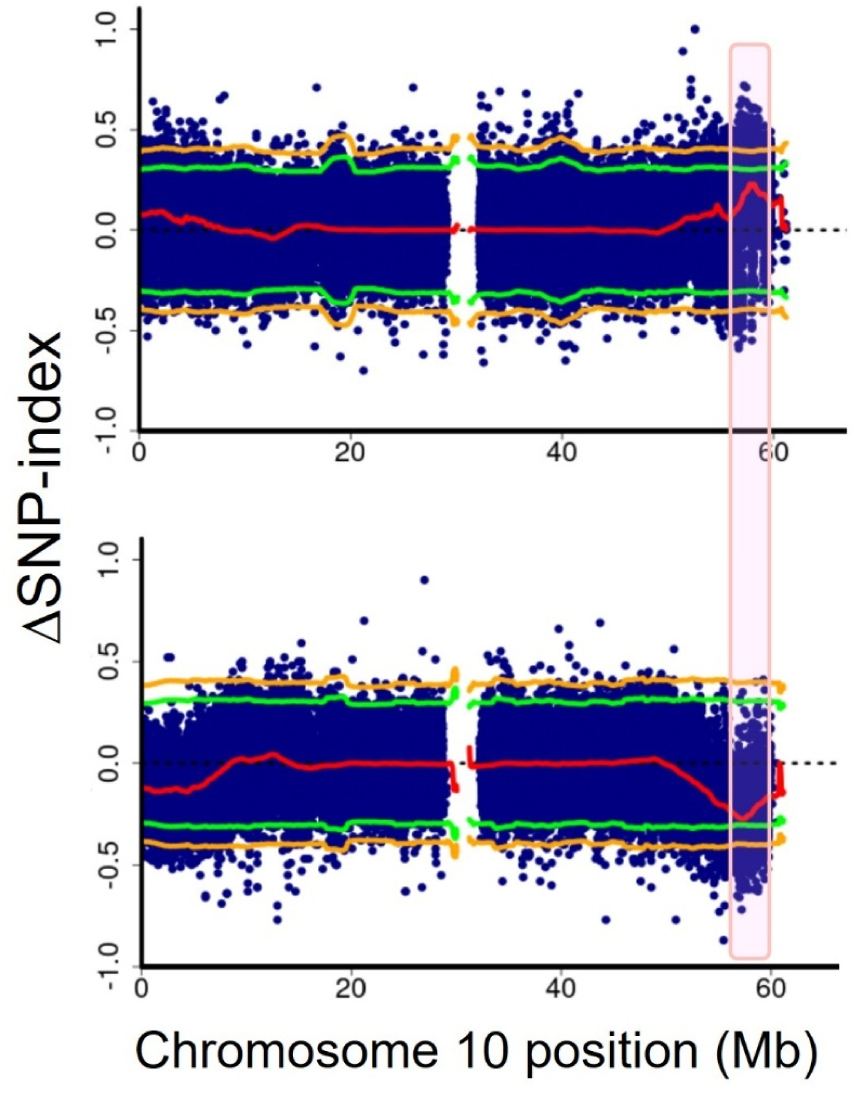
A resistance QTL at the distal end of chromosome 10. QTL map on chromosome 10 based on the resistant (upper) and the susceptible (lower) parental genomes as the internal reference. Blue dot shows the vertical position of a SNP and the red line along the x-axis shows the average ΔSNP-index. The green and orange lines show statistical thresholds (p-value = 0.05 and 0.01). The peak shows the QTL region (p-value approximately 0.05). The QTL harbors a BTB/POZ candidate gene.

**Fig. S7.**
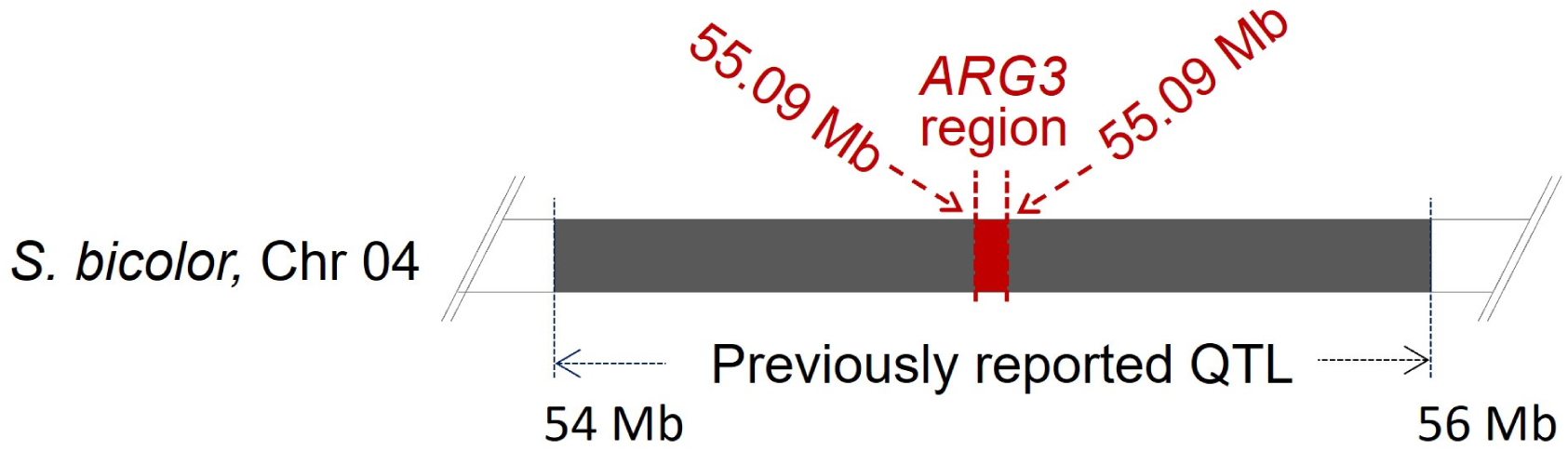
*ARG3* was located at the center of a previously reported anthracnose resistance QTL in IS18760. *ARG3* was located at the center of a previously reported anthracnose resistance QTL in IS18760. In the current study, the 30 kb *ARG3* locus (55.09-55.12 Mb) in IS18760 was identified independently of earlier findings, yet it resides within the previously reported 2 Mb (54-56 Mb).

## Supplementary File 1. Aligned coding sequences of the tyrosine kinase domain gene (TK) in the mapping interval in parental lines and the other variant line RTx430

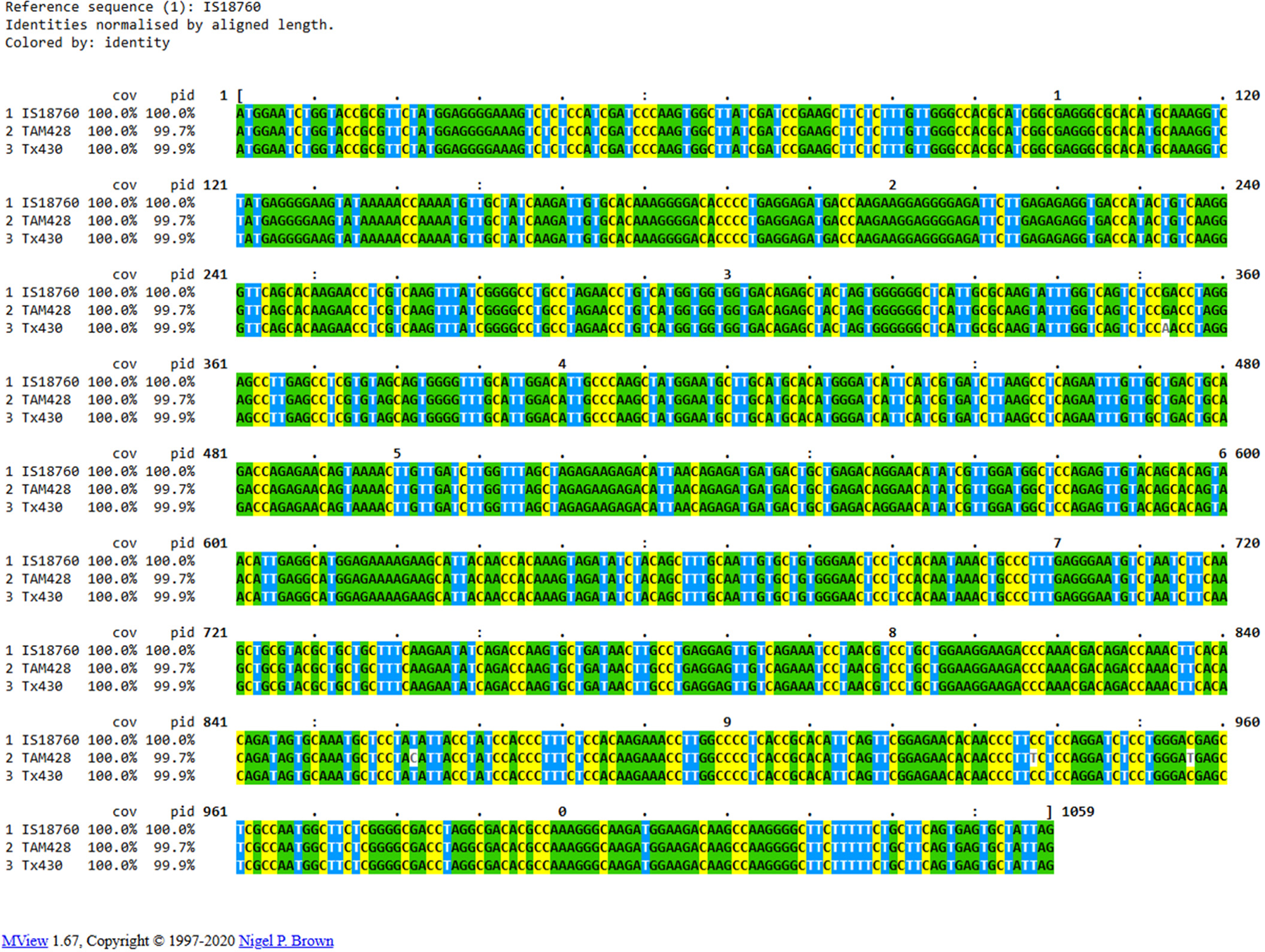

## Supplementary File 2. *ARG3* candidate transcript sequence in 18 *Sorghum* lines and subspecies that represent the 111 pangenome lines aligned with parental lines

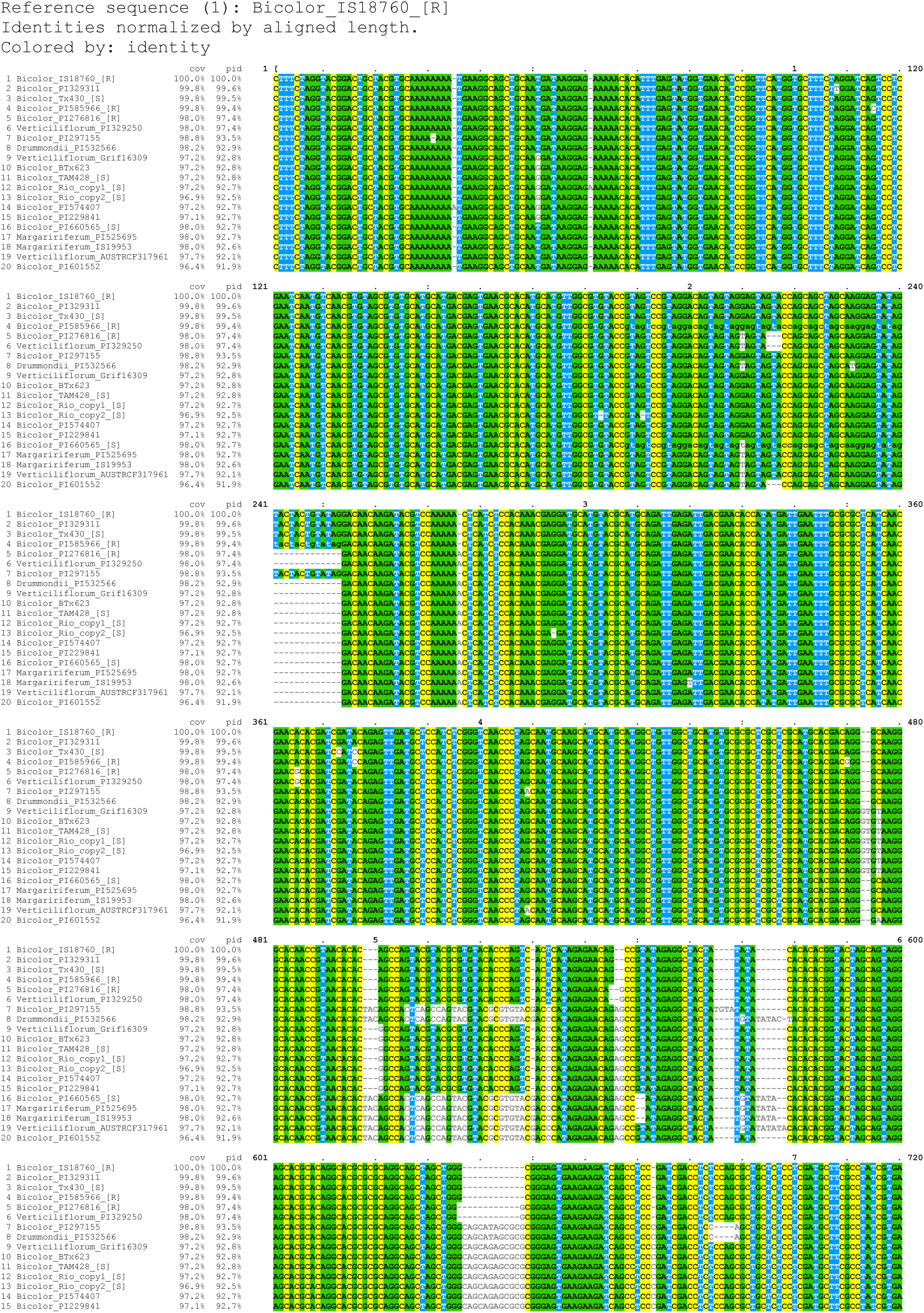
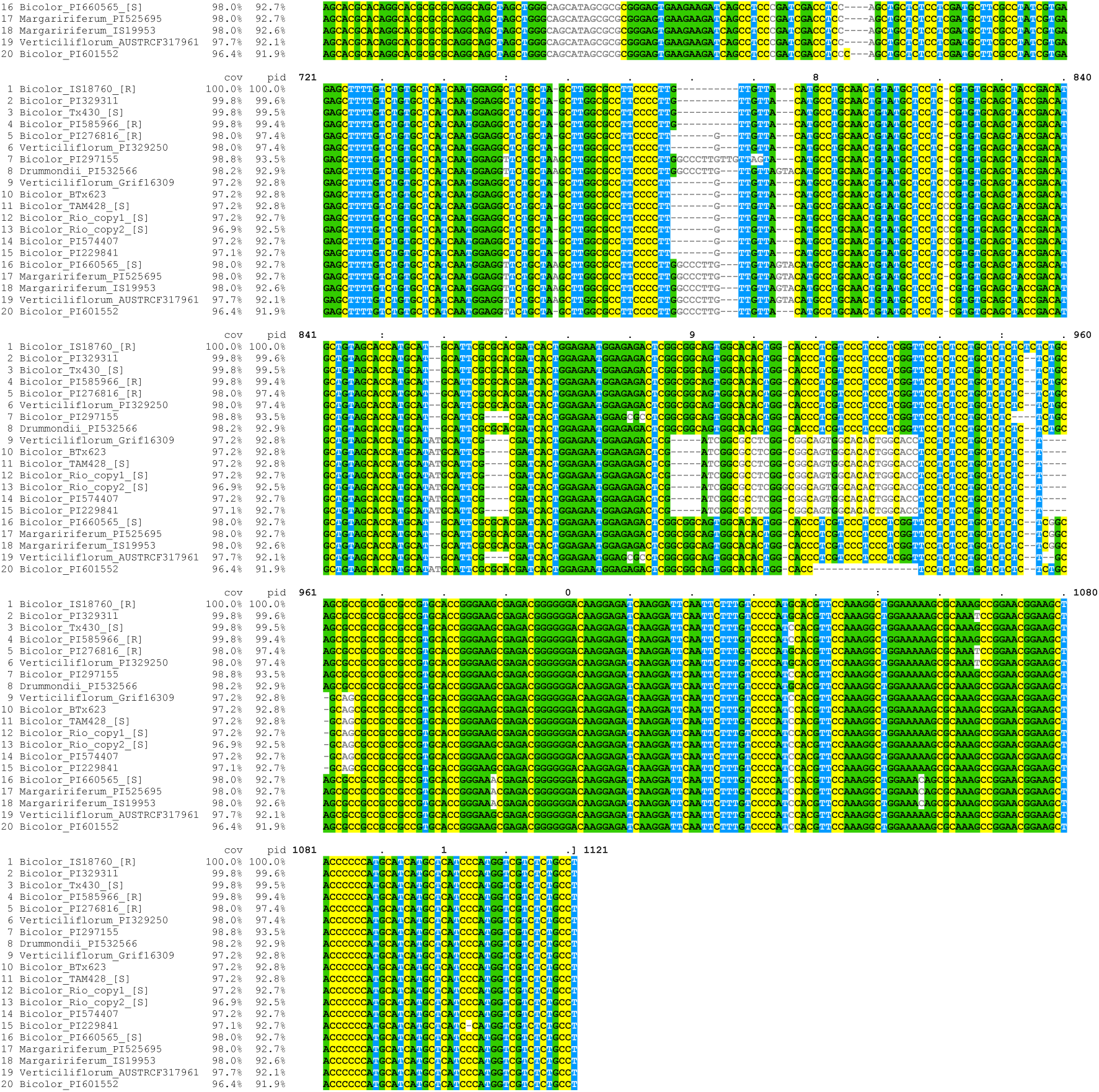

